# Immuno-informatics Characterization SARS-CoV-2 Spike Glycoprotein for Prioritization of Epitope based Multivalent Peptide Vaccine

**DOI:** 10.1101/2020.04.05.026005

**Authors:** Saba Ismail, Sajjad Ahmad, Syed Sikander Azam

## Abstract

The COVID-19 pandemic caused by SARS-CoV-2 is a public-health emergency of international concern and thus calling for the development of safe and effective therapeutics and prophylactics particularly a vaccine to protect against the infection. SARS-CoV-2 spike glycoprotein is an attractive candidate for vaccine, antibodies and inhibitor development because of many roles it plays in attachment, fusion and entry into the host cell. In this study, we characterized the SARS-CoV-2 spike glycoprotein by immune-informatics techniques to put forward potential B and T cell epitopes, followed by the use of epitopes in construction of a multi-epitope peptide vaccine construct (MEPVC). The MEPVC revealed robust host immune system simulation with high production of immunoglobulins, cytokines and interleukins. Stable conformation of the MEPVC with a representative innate immune TLR3 receptor was observed involving strong hydrophobic and hydrophilic chemical interactions, along with enhanced contribution from salt-bridges towards inter-molecular stability. Molecular dynamics simulation in solution aided further in interpreting strong affinity of the MEPVC for TLR3. This stability is the attribute of several vital residues from both TLR3 and MEPVC as shown by radial distribution function (RDF) and a novel analytical tool axial frequency distribution (AFD). Comprehensive binding free energies estimation was provided at the end that concluded major domination by electrostatic and minor from van der Waals. Summing all, the designed MEPVC has tremendous potential of providing protective immunity against COVID-19 and thus has the potential to be considered in experimental studies.

## 1. Introduction

In December 2019, a new strain of coronavirus emerged in Wuhan city of Hubei province in China and has since spread globally. The virus belongs to clade B of family Coronaviridae in the order Nidovirales, and genera Betacoronavirus and caused pulmonary disease outbreak [1,2]. It is positive-sense RNA, enveloped and non-segmented virus and named as SARS-CoV-2 as it share 82% sequence identity with SARS coronavirus (SARS-CoV)[3,4]. SARS-CoV-2 caused coronavirus disease-19 (COVID-19) and evidence suggest a zoonotic origin of this disease [5]. Though the zoonotic transmission is not completely understood but facts provide the ground that it proliferates from the seafood market Huanan in Wuhan and human-to-human transmission resultant into the exponential increase in number of cases [6,7]. As of March 24, 386,332 cases are reported worldwide with 16,747 deaths and 102,333 recovered patients. Among the active cases, 267,252 are currently infected, 255,166 (95%) are in mild conditions and 12,086 (5%) are seriously ill. serisouly illed. Among the 119,080 closed cases, 102,333 (86%) are recovered whereas 16,747 (14%) die. On March 11, the World Health Organization (WHO) affirmed COVID-19 as a pandemic (https://www.worldometers.info/coronavirus/).

SARS-CoV-2 utilizes a highly glycosylated, homotrimeric class I viral fusion spike protein to enter into host cells [8]. This protein is found in a metastable pre-fusion state which undergoes a structural rearrangement facilitating viral membrane fusion with the host cell[9–11]. The binding of S1 subunit to a host-angiotensin-converting enzyme initiates this process and disrupts the prefusion trimeric structure resulting into S1 subunit dispersion and stabilizes the S2 subunit to a post-fusion conformation[12]. The receptor-binding domain (RBD) of S1 goes through a hinge-like conformational change that temporarily hides or exposes the determinants of receptor binding in order to occupy a host-cell receptor [11]. Down and up conformation states are recognized where former is related to the receptor-inaccessible state and the later one explains receptor-accessible state and considered as less stable [13–16]. This critical role of the spike protein makes it an important target for antibody-mediated neutralization, and detailed study of the pre-fusion S structure would provide information at atomic-level helping in the design and development of a vaccine [17–21]. Current data indicates that SARS-CoV-2 spike and SARS-CoV spike both share the same functional receptor (host cell) —angiotensin-converting enzyme 2 (ACE2) [22,23]. Interestingly, ACE2 binds to SARS-CoV-2 spike ectodomain with ~15 nM affinity, about 10-20 folds higher than ACE2 binding to SARS-CoV spike [24]. One possible reason for SARS-CoV-2 capability of spreading infection from human-to-human is SARS-CoV-2 spike’s high affinity for human ACE2[25]. Series of cellular immune and humoral responses can be triggered by SARS-CoV-2 infections [26]. Immunoglobulin G (IgG) and IgM were noticeable after the 2 weeks of onset of infection which are specific antibodies to SARS-CoV-2. High titers of neutralizing antibodies and SARS-CoV-2 specific cytotoxic T lymphocyte responses have been identified in the patients who have improved from SARS-CoV-2. This phenomenon clearly suggest that both cellular and humoral immune reactions are vital to clearing the SARS-CoV-2 infection [26–30].

The study presented, herein, is an attempt to get insights about antigenic determinants of SARS-CoV-2 spike glycoprotein and highlight all antigenic epitopes [31] of the spike that can be used specifically for the design of a multi-epitope peptide vaccine construct (MEPVC) [32]to counter COVID-19 infections. The epitopes predicted by immunoinformatics techniques were fused together as well as to β-defensin adjuvant [33,34] to boost the antibody production and long-lasting immunological responses. Blind docking protocol was implemented to gain MEPVC best possible binding mode with respect to representative innate immune receptor TLR3 [35]. The docked solution was further utilized in dynamics simulations to study the structural dynamics and stability of the complex [36]. Finally, to confirm intermolecular interactions and point all noteworthy residues crucial for intermolecular stability, binding free energies [37] for the system were estimated. In a nutshell, this study reflects excellent outcomes for experimental vaccinologists to develop a vaccine to control pandemic COVID-19 infections.

## 2. Materials and Methods

Complete flow of the *in silico* designed methodology for MEPVC targeting SARS-CoV-2 9 spike is demonstrated in **Fig. 1.**

**Fig.1.**
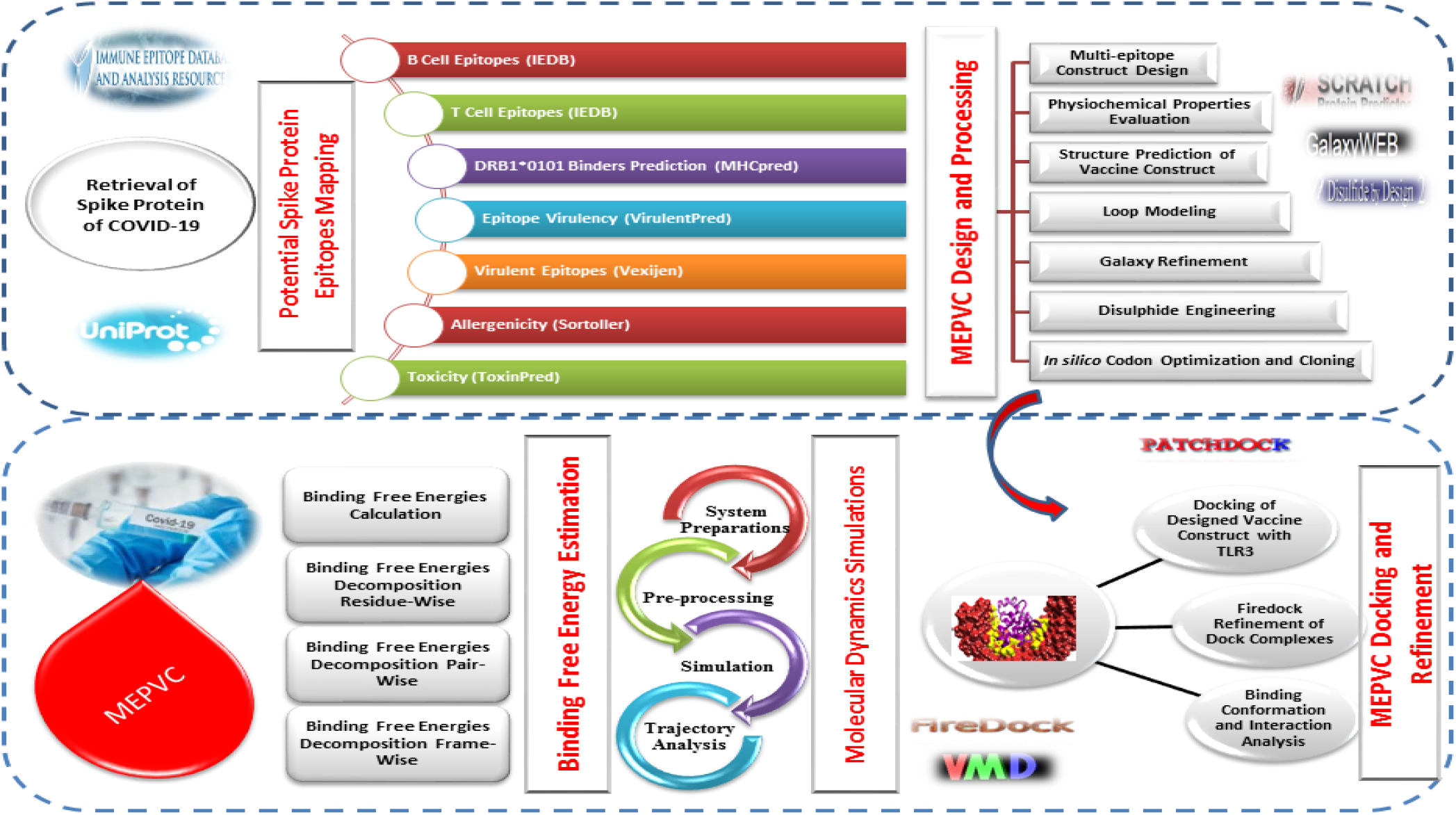
Computational approach adopted for the design of a SARS-CoV-2 spike protein based MEPVC.

### 2.1. Epitopes Mapping for Spike Protein

The amino acid sequence of spike glycoprotein protein from SARS-CoV-2 was retrieved from NCBI SARS-CoV-2 data hub and considered first in the epitope mapping phase, where T cell epitopes derived from B-cell were predicted using immune epitopes data base (IEDB)[38]. Linear B cell epitopes were mapped using Bepipred Linear Epitope Prediction 2.0 [39] and those with score greater than 0.5 were subjected to T-cell epitopes identification step. The epitopes were projected for association with reference set of major histocompatibility complex (MHC): MHC class I [40] and MHC class II [41] alleles on the basis of percentile score. Epitopes with lowest percentile score were considered only as high affinity binders. The selected epitopes were then used in MHCPred 2.0 [42] to decipher their binding affinity potential for predominant HLA II DRB*0101 and only those with IC_50_ value < 100 nM were categorized as excellent DRB*0101 binders [43]. VirulentPred [44] was employed next to reveal virulent nature of the epitopes setting the cut-off to 0.5. Antigenic epitopes were highlighted by VaxiJen 2.0 [45]. Allergenic epitopes were discarded through AllerTop 2.0 [46] and toxic potential of non-allergic epitopes was evaluated via ToxinPred [47]. The non-toxic epitopes were lastly investigated for their ability of inducing IFN-γ using an IFN epitope server [48]. Conservation across the world population of the final set of epitopes was done through IEDB epitope conservation analysis tool [49].

### 2.2. Multi-Epitopes Peptide Vaccine Construct (MEPVC) Designing and Post Analysis

All filtered epitopes were linked together through AAY linkers [50] to design a multi-epitope peptide (MEP). The resultant peptide was further linked to an immunological β-defensin (an adjuvant) to construct a multi-epitope peptide vaccine construct (MEPVC) and in this way immunogenicity can be enhanced. The physicochemical properties of designed MEPVC was predicted by ProtParam tool [51] of EXPASSY server. The three dimensional (3D) structure of the MEPVC was modeled *ab initio* by 3Dpro of SCRATCH protein server [52]. Following, loop modeling was done in the 3D structure of MEPVC via GlaxyLoop [53] from GlaxyWeb and subsequently refined through GalaxyRefine [54]. Disulfide engineering was applied on the MEPVC refined model via Design 2.0 [55] as disulfide bonds strengthen structure stability. The MEPVC sequence was translated reversibly for optimization of codon usage according to *Escherichia coli* K12 expression system in order to get high expression rate [56]. For this, Java Codon Adaptation Tool (JCat) [57] was used and expression of the cloned MEPVC was assessed by GC *%* and codon adaptation index (CAI) value. SnapGene (https://www.snapgene.com/) was used to clone the optimized MEPVC cDNA into pET-28a (+) expression vector.

### 2.3. *In Silico* Immune Profiling of MEPVC

Immunogenic potential of the MEPVC was conducted *in silico* using C-ImmSim server [58,59]. The server used machine learning techniques along with position-specific scoring matrix (PSSM) for prediction of the host immune system response to the antigen. The immune system responds from three mammalian anatomical regions: bone marrow, lymph nodes and thymus. The input parameters for the immune simulations are as follows: number of steps (100), volume (10), random seed (12345), HLA (A0101, A0101, B0702, B0702, DRB1_0101, DRB1_0101), number of injection set to 1. The rest of parameters were treated as default.

### 2.4. Molecular Docking of MEPVC

The MEPVC affinity for an appropriate immune receptor as an agonist was checked in the step of molecular docking[60]. TLR3 available under PDB id of 1ZIW was retrieved and used as a receptor molecule. TLR3 also named as CD283 is a transmembrane protein belongs to the family of pattern recognition receptor [61]. It detects viral infection-associated dsRNA and evoke the activation of interferon regulatory transcription factor (IRF3) and (Nuclear Factor kappa-light-chain-enhancer) NF-kB [62]. Unlike other TLRs, TLR3 uses TIR-domain-containing adapter-inducing interferon-β (TRIF) as a primary adapter [63]. IRF3 eventually induces the development of type I interferons leading to the activation of innate immune system and eventually to long lasting adaptive immunity [64]. The TLR3 receptor and MEPVC were used in a blind docking approach through an online PATCHDOCK server interface [65]. The interacting molecules were docked according to the shape complementarity principle. The clustering RMSD is allowed to default 4.0 Å. The output docked solutions were immediately refined with Fast Interaction Refinement in Molecular Docking (FireDock) server [66] which provides an efficient framework for refining PATCHDOCK complexes. The refined complexes were examined and one with lowest global energy was considered as top ranked. The opted complex was subjected to in-depth MEPVC conformation with respect to the TLR3 using UCSF Chimera 1.13.1 [67].

### 2.5. Molecular Dynamics (MD) Simulation

MD simulation was applied on the selected top complex for 50-ns to understand complex dynamics and stability for practical applications. This assay was categorized into three phases: (i) parameters file preparation (ii) pre-processing, and (iii) simulation production [68]. In first phase using an antechamber module of AMBER16 [69], complex libraries and set of parameters for TLR3 and MEPVC were generated. The complex system was solvated into 12 Å TIP3P solvation box achieved through Leap module of AMBER. The intermolecular and intramolecular interactions of the system were determined by ff14SB force [70] field. Counter ions in the form of Na^+^ were added to the system for charge neutralization. In the system pre-processing stage, complex energy was optimized through several rounds: minimization of complete set of hydrogen atoms for 500 steps, minimization of system solvation box energy for 1000 steps with restraint of 200 kcal/mol – Å^2^ on the remaining system, minimization of complete set of system atoms again for 1000 steps with applied restraint of 5 kcal/mol −Å^2^ applied on system carbon alpha atoms, and 300 steps of minimization on system non-heavy atoms with restraint of 100 kcal/mol −Å^2^ on other system components. The complex system then underwent a heating step where the complex was heated gradually to 300K through NVT ensemble, maintained through Langevin dynamics [71] and SHAKE algorithm [72] to restrain hydrogen bonds. Complex equilibration was achieved for 100-ps. Pressure on the system was maintained using NPT ensemble allowing restraint on Cα atoms of 5 kcal/mol – Å^2^. In the simulation production, trajectories of 50-ns were produced on time scale of 2-fs. Non-bounded interactions were differentiated by describing cut-off distance of 8.0 Å.CPPTRAJ module [73] was lastly used for statistical computation of different structure parameters to probe complex stability. The MD simulation trajectories were visualized and analyzed in Visual Molecular Dynamics (VMD) 1.9.3 [74].

### 2.6. Axial Frequency Distribution Analysis

Axial frequency distribution or simply AFD [75] is a novel analytical technique run on simulation trajectories to access ligand 3D conformation with respect to reference receptor atom. Such local structural movements are not captured by any other available technique. AFD can be mathematically presented by Eq.I,

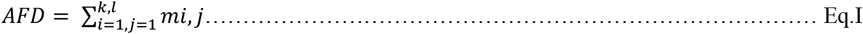

where, *i* and *j* are ligand atom coordinates on X and Y axis with cut-off value *k* and *l*, respectively. The *mi,j* sums interactions frequency that fall in the coordinate (*i,j*).

### 2.7. Calculating Binding Free Energy for TLR3-MEPVC

The interaction energy and solvation free energy for TLR3 receptor, MEPVC, TLR3-MEPVC complex were calculated utilizing the MMPBSA.py module [76] of AMBER16. Also, an average of the above was estimated as a net binding free energy of the system. The binding free energy was computed through MM-PBSA method and its counterpart MM-GBSA of AMBER with objective to derive the difference between bound and unbound states of solvated conformations of the same molecule [77]. Mathematically, the binding free energy can be calculated though Eq.II,

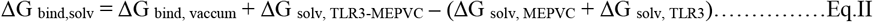

For all three states of the system, the solvation energy was calculated by solving either Poisson Boltzman (PB) or Generalized Born (GB) equation and thus it will give electrostatic contribution of the solvation state. It also allow the addition of empirical term for hydrophobic contributions as shown in Eq.III.

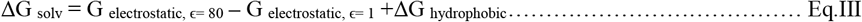

Calculation of the average interaction energy between TLR3 and MEPVC gives to delta-ΔG vacuum (Eq.IV)

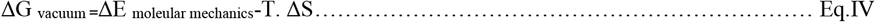

## 3. Results and Discussion

### 3.1. Prioritization of Potential Epitopes

The SARS-CoV-2 spike protein was targeted for MEPVC designing because of many filters it fulfilled required for a potential vaccine candidate. First, it does not share any significant homology to the human host and as such chances of autoimmune responses are negligible [78]. Second, the protein is also not found to have any sequence identity to the mouse proteome and thus accurate immunological findings can be deciphered from *in vivo* mice experimentations [79]. This spike protein only harbored one transmembrane helix ensuring the wet lab protein cloning and expression for antigen analysis easy [80]. Antigenicity is another factor that make this candidate highly suitable for vaccine designing as this allows efficient binding to the products of host immune system [81]. Further, this protein is strongly adhesive which makes it an excellent target for creation of adhesion based vaccine [82]. Lastly, all the sequences of SARS-CoV-2 spike protein are highly conserved thus a vaccine based on its sequence will be highly likely to have broad spectrum immunological implications [83]. Prioritization of potential epitopes for the SARS-CoV-2 spike protein commenced with the mapping of B-cell epitopes that predicted total of 34 epitopes of vary length ranging from one to 62 (**S Table 1**). The average score predicted for these B cell epitopes is 0.470, maximum (max) of 0.696 and minimum (min) of 0.188 (**Fig.2**). Each B-cell epitope was then analyzed in MHC-1 alleles binding regions prediction [84]. A set of reference alleles to which these epitopes interact with are: HLA-A*01:01, HLA-A*02:01, HLA-A*02:01, HLA-A*02:03, HLA-A*02:03, HLA-A*02:06, HLA-A*02:06, HLA-A*03:01, HLA-A*03:01, HLA-A*11:01, HLA-A*11:01, HLA-A*23:01, HLA-B*08:01, HLA-A*23:01, HLA-A*24:02, HLA-A*24:02, HLA-A*26:01, HLA-A*26:01, HLA-A*30:01, HLA-A*30:01, HLA-B*57:01, HLA-A*30:02, HLA-A*31:01, HLA-B*58:01, HLA-A*32:01, HLA-A*33:01, HLA-A*33:01, HLA-A*68:01, HLA-A*68:01, HLA-A*68:02, HLA-A*68:02, HLA-A*30:02, HLA-B*07:02, HLA-B*51:01, HLA-B*07:02, HLA-B*08:01, HLA-B*15:01, HLA-B*15:01, HLA-B*35:01, HLA-A*31:01, HLA-B*35:01, HLA-B*40:01, HLA-B*40:01, HLA-B*44:02, HLA-B*44:02, HLA-B*44:03, HLA-B*44:03, HLA-B*51:01, HLA-A*01:01, HLA-B*53:01, HLA-B*53:01, HLA-B*57:01, HLA-A*32:01, and HLA-B*58:01. These alleles have > 97% population coverage. The predicted epitopes were then screened and stringent criteria of lowest percentile score was used to choose the excellent binders. Afterward, the B cell epitopes were simultaneously run in MHC-II alleles binding [85]. Likewise MHC-I, reference set of MHC-II binding were: HLA-DRB4*01:01, HLA-DRB1*04:01, HLA-DRB1*04:05, HLA-DRB1*07:01, HLA-DRB1*09:01, HLA-DRB1*11:01, HLA-DRB1*03:01, HLA-DRB1*13:02, HLA-DRB1*15:01, HLA-DRB3*01:01, HLA-DRB1*12:01, HLA-DRB3*02:02, HLA-DRB1*08:02, HLA-DRB1*01:01, and HLA-DRB5*01:01. The MHC-II predicted epitopes were also filtered on basis of percentile score and then cross checked with the selected MHC-I allele and those common in both classes were considered only which were 50 in numbers. The shortlisted common MHC-I and MHC-II epitopes then subjected to antigenicity check. In this check, ability of the filtered B-cell derived T-cell epitopes ability to evoke and bind to products of adaptive immunity. This yielded 38 epitopes all of which have strong ability to bind to the most prevalent DRB*0101 with average IC50 score of 35.6552, max of 98 and min of 0.89.The antigenic epitopes then underwent allergenicity check to discard allergic peptides that may cause allergic reactions [86]. This resulted into 31 epitopes. Non-toxic epitopes were 7 whereas 6 were IFN-gamma producer (**Fig.3**). The set of epitopes obtained at different stages of epitope mapping phase is tabulated in **Table 1**. These epitopes appear to provide coverage to 98% of the world population (**Fig.4**). The coverage can split as: 98.54% in East Asia, 96.06% in Northeast Asia, 97.46% in South Asia, 95.83% in Southeast Asia, 94.29% in Southwest Asia, 98.53% in Europe, 92.81% in East Africa, 89.82% in Central Africa, 96.45% in West Africa, 97.86% in North Africa, 94.81% in South Africa, 99.09% in West Indies, 98.74% in North America, 25.01% in Northern America, 90.91% in South America, and 95.58% in Oceania.

**Fig.2.**
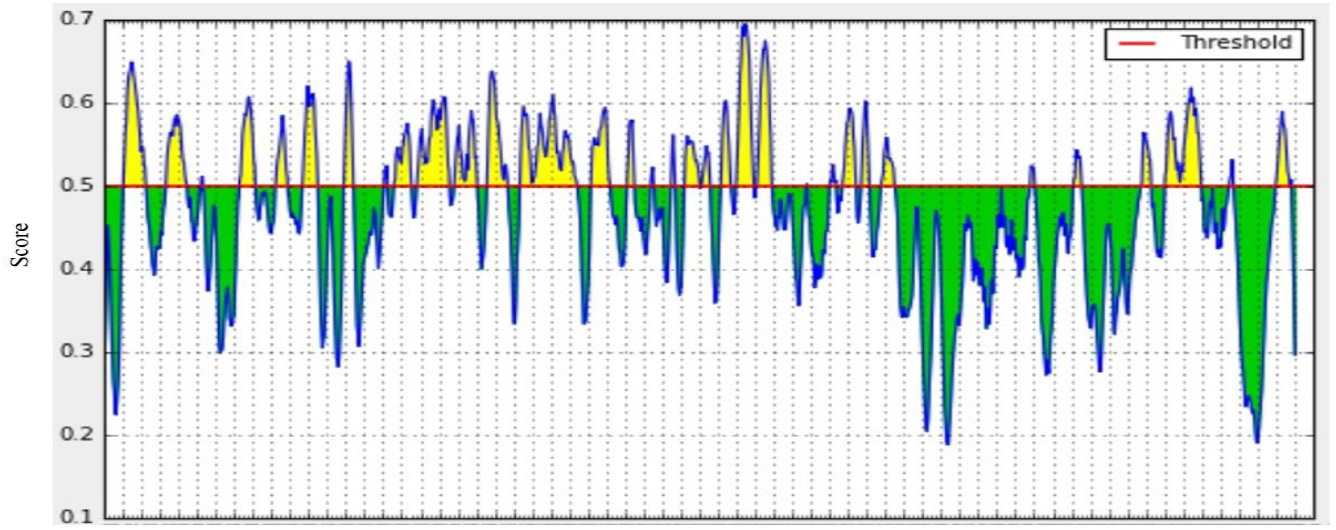
B cell epitopes prediction by IEDB Bepipred linear epitope prediction method. Yellow and green peaks are those predicted as epitopes and non-epitopes, respectively.

**Fig.3.**
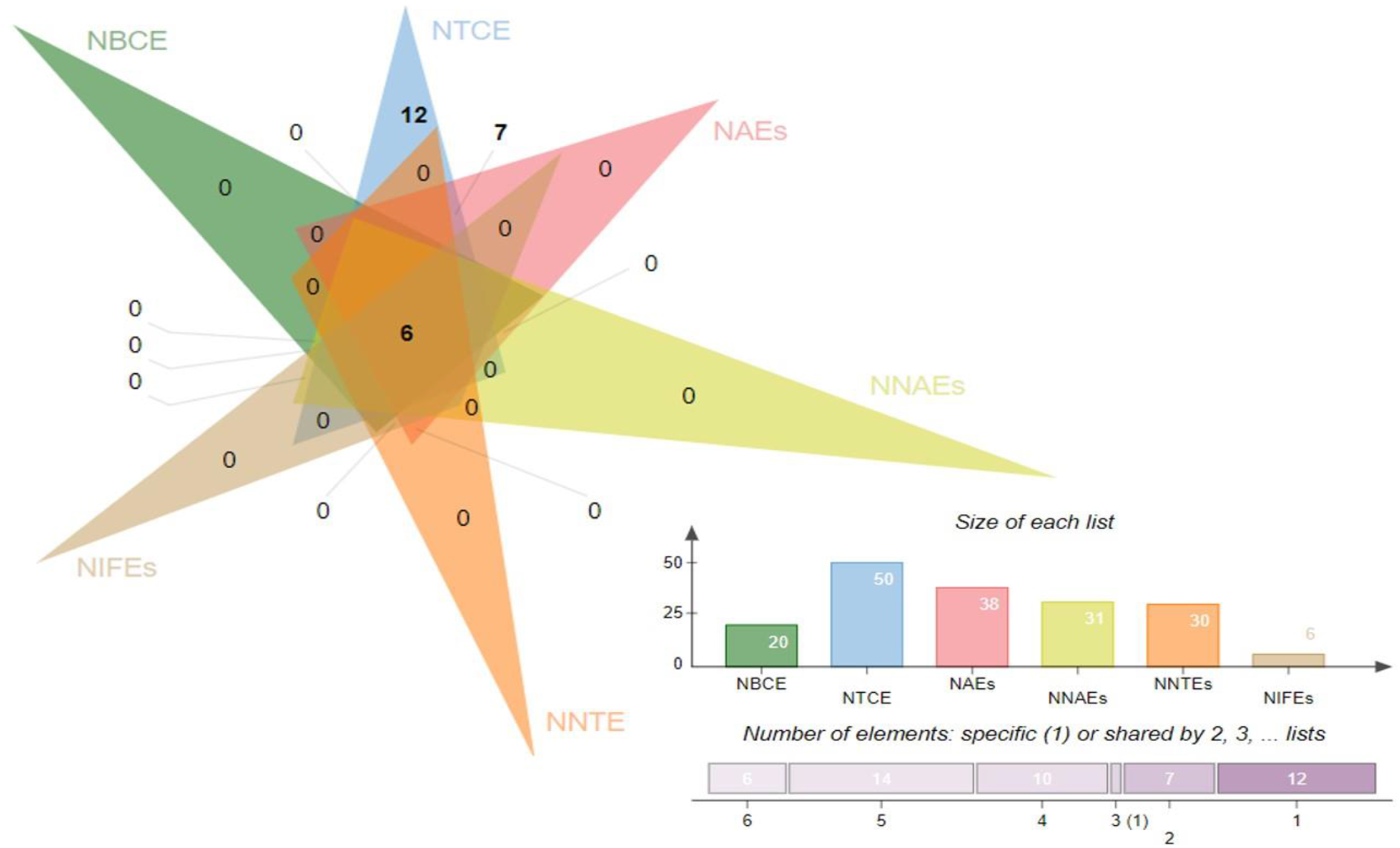
Venn diagram for the number of different categories of epitopes predicted at epitope mapping stage. NBCE (number of B cell epitopes), NTCE (number of T cell epitopes), NAEs (number of antigenic epitopes), NNAES (number of non-allergic epitopes), NNTEs (number of non-toxic epitopes), and NIFEs (number of IFN-gamma epitopes).

**Fig.4.**
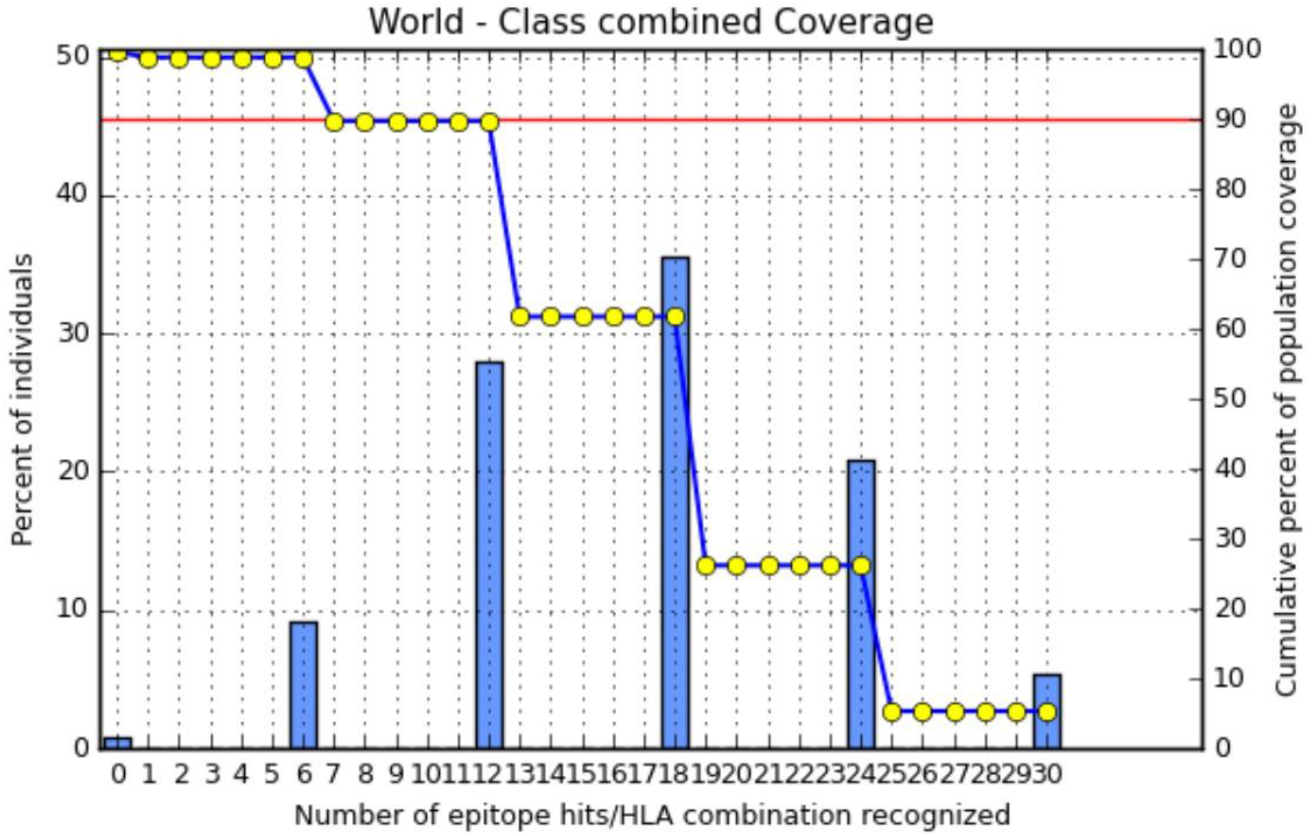
World population coverage of the final set of predicted epitopes.

**Table 1.** Number of epitopes obtained at each step of epitope mapping phase.

| B- Cell Epitopes | T-Cell Epitopes | Percentile score |  | MHC Pred Score (nM) | Antigenicity | Allergenicity | Toxicity | IFN-gamma producer | Final Selection |
| --- | --- | --- | --- | --- | --- | --- | --- | --- | --- |
|  |  | MHC 1 | MHC II |  |  |  |  |  |  |
| SQCVNLTRTQLPPAYTNSFTRGVY | TRTQLPPAY | 1.04 | 34 | 12.91 | 1.2923 | Non-Allergen | Non-Toxin |  | - |
|  | RTQLPPAYT | 1.4 | 34 | 76.31 | 1.2195 | Non-Allergen | Non-Toxin |  | - |
|  | TTRTQLPPA | 1.4 | 31 | 10.4 | 1.254 | Non-Allergen | Non-Toxin |  | - |
| FSNVTWFHAIHVSGTNGTKRFDN | FSNVTWFHA | 1.4 | 1.5 | 3.41 | 0.8156 | Non-Allergen | Non-Toxin |  | - |
|  | TWFHAIHVS | 6.6 | 5.6 | 48.53 | 0.5374 | Non-Allergen | - |  | - |
| DPFLGVYYHKNNKSWME | VYYHKNNKS | 9.8 | 2.1 | 26.3 | 0.451 | Allergen | - |  | - |
|  | YYHKNNKSW | 0.36 | 2.1 | 80.17 | 0.4536 | Non-Allergen | Non-Toxin |  | - |
| MDLEGKQGNFKNL | KQGNFKNLR | 2.65 | 7.1 | 31.7 | 0.9558 | Non-Allergen | Non-Toxin |  | - |
| KHTPINLVRDLPQGFS | KHTPINLVR | 3.2 | 0.51 | 34.36 | 1.0552 | Non-Allergen | Non-Toxin |  | - |
| TPGDSSSGWTA | PGDSSSGWT | 31 | 66 | 98 | 0.1337 | Non-Allergen | Non-Toxin |  | - |
| KSFTVEKGIYQTSNFRVQP | YQTSNFRVQ | 1.95 | 3.92 | 5.19 | 0.7821 | Non-Allergen | Non-Toxin |  | - |
| FPNITNLCPFGEVFNATRFASVYAWNRRKRISNCVA | PNITNLCPF | 0.06 | 0.09 | 9.18 | 1.6189 | Non-Allergen | Non-Toxin |  | - |
|  | NITNLCPFG | 15 | 2.7 | 19.5 | 1.5251 | Allergen | Non-Toxin |  | - |
|  | YAWNRRKRIS | 4.45 | 0.52 | 9.2 | 0.8209 | Non-Allergen | Non-Toxin | Positive | Selected |
|  | FNATRFASV | 0.2 | 2.6 | 24.6 | 0.5609 | Non-Allergen | Non-Toxin |  | - |
|  | FPNITNLCP | 0.06 | 0.09 | 44.87 | 1.6A218 | Non-Allergen | Non-Toxin |  | - |
| YNSASFSTFKCYGVSPTKLNDLCFT | FSTFKCYGV | 0.4 | 9.5 | 8.45 | 0.414 | Non-Allergen | Non-Toxin |  | - |
|  | PTKLNDLCF | 0.52 | 32 | 12.85 | 2.5304 | Non-Allergen | Non-Toxin |  | - |
| GDEV RQIAPGQTGKIADYNYKLP | GQTGKIADY | 0.77 | 15.5 | 12.11 | 1.4019 | Non-Allergen | Non-Toxin |  | - |
|  | IAPGQTGKI | 1.4 | 30.33 | 66.99 | 1.6527 | Non-Allergen | Non-Toxin | Positive | Selected |
| NLDSKVGGNYNLYRLFRKSNLKPFERDISTEYIYQAGSTPCNGV<br>EGFNCFYPLQSYGFQPTN | FRKSNLKP | 2.6 | 3.1 | 3.41 | 0.628 | Non-Allergen | Non-Toxin |  | - |
|  | NLKPFERDI | 7.6 | 1.8 | 6.41 | 0.9116 | Non-Allergen | Non-Toxin |  | - |
|  | KSNLKPFER | 0.14 | 1.9 | 38.9 | 0.949 | Non-Allergen | Non-Toxin |  | - |
| ELLHAPATVCGPKKSTNLVKN | PKKSTNLVK | 3.3 | 17 | 85.11 | 0.4877 | Non-Allergen | Non-Toxin |  | - |
|  | KKSTNLVKN | 1.55 | 14 | 90.7 | 0.4876 | Non-Allergen | Non-Toxin |  | - |
| NCTEVPVAIHADQLTPT | NCTEVPVAI | 3.8 | 23.5 | 40.18 | 0.5598 | Allergen | Non-Toxin |  | - |
| RVYSTGNSNVFQ | VYSTGNSNVF | 0.17 | 6.2 | 55 | 0.3099 | Allergen | Non-Toxin |  | - |
| VNNSYECDIPI | NSYECDIPI | 1.9 | 8.4 | 23.93 | 0.2216 | Non-Allergen | - |  | - |
| ASYQTQTNSPRRARSVASQ | PRRARSVAS | 0.17 | 3.9 | 13.3 | 0.4829 | Non-Allergen | Non-Toxin | Positive | Selected |
|  | ASYQTQINS | 4.9 | 8.6 | 23.88 | 0.5213 | Non-Allergen | Non-Toxin |  | - |
|  | SPRRARVA | 0.1 | 4 | 0.89 | 0.7729 | Non-Allergen | Non-Toxin |  | - |
| YTMSLGAENSVAYSNN | MSLGAENSV | 1.45 | 3.3 | 60.95 | 0.873 | Non-Allergen | Non-Toxin |  | - |
|  | SLGAENSV | 0.67 | 4.9 | 61.5 | 0.6459 | Allergen | Non-Toxin |  | - |
| RNFYEPQIITTD | YEPQIITTD | 3 | 2.8 | 47.86 | 0.8297 | Allergen | Non-Toxin |  | - |
|  | NFYEPQIIT | 11.35 | 17 | 85.11 | 0.1785 | Allergen | Non-Toxin |  | - |
| VNNTVYDPLQPELDSFKEELDKYFKNHTSPDVLGDISGI | TVYDPLQPE | 2.2 | 33 | 5.448 | 0.5391 | Non-Allergen | Non-Toxin | Positive | Selected |
|  | VYDPLQPEL | 2.91 | 33.3 | 14.79 | 0.4525 | Non-Allergen | Non-Toxin | Positive | Selected |
|  | PDVDLGDIS | 7.75 | 30 | 20.65 | 2.0111 | Non-Allergen | Non-Toxin |  | - |
|  | FKNHTSPDV | 6.4 | 2.2 | 45.71 | 0.4846 | Non-Allergen | Non-Toxin | Positive | Selected |
| SCCKFEDEDDSEPVLKG | HTSPDVLG | 2.9 | 30 | 67.45 | 1.5209 | Non-Allergen | Non-Toxin |  |  |

### 3.2. MEPVC Designing

Prioritized T cell epitopes derived from B cells were fused together tandemly by AAY linkers to make a multi-epitope peptide (MEP). AAY linker avoid formation of junctional epitopes and as such enhance epitope presentation [87]. To the N-terminal of MEP, EAAAK linker was added to attach β-defensin as an adjuvant leading to the design of a MEPVC. The MEPVC is schematically shown in **Fig.5A**. MEPVC offers many advantages compared to a separate antigenic peptide. Such vaccines induce both CD4+ and CD8+ responses and the antigens optimization are optimal. EAAAK is a rigid spacer and allow separation of the attached domain and promoting efficient immune processing of the epitopes [88]. β-defensins are potent immune adjuvants as they are capable of significantly enhancing production of lymphokines resulting into antigen-specific Ig production and T cell-dependent cellular immunity. The sequence of MEPVC is: GIINTLCKYYCRVRGGRCCVCSCCPKEEQIGKCSTRGRKCCRRKKECAAKYAWNRKCISACYIAPGQTGKICCYPRRARSVCSACYTVYDPCQPCAAYVYDPLCPELCAYCKN HTSCDV.

**Fig.5.**
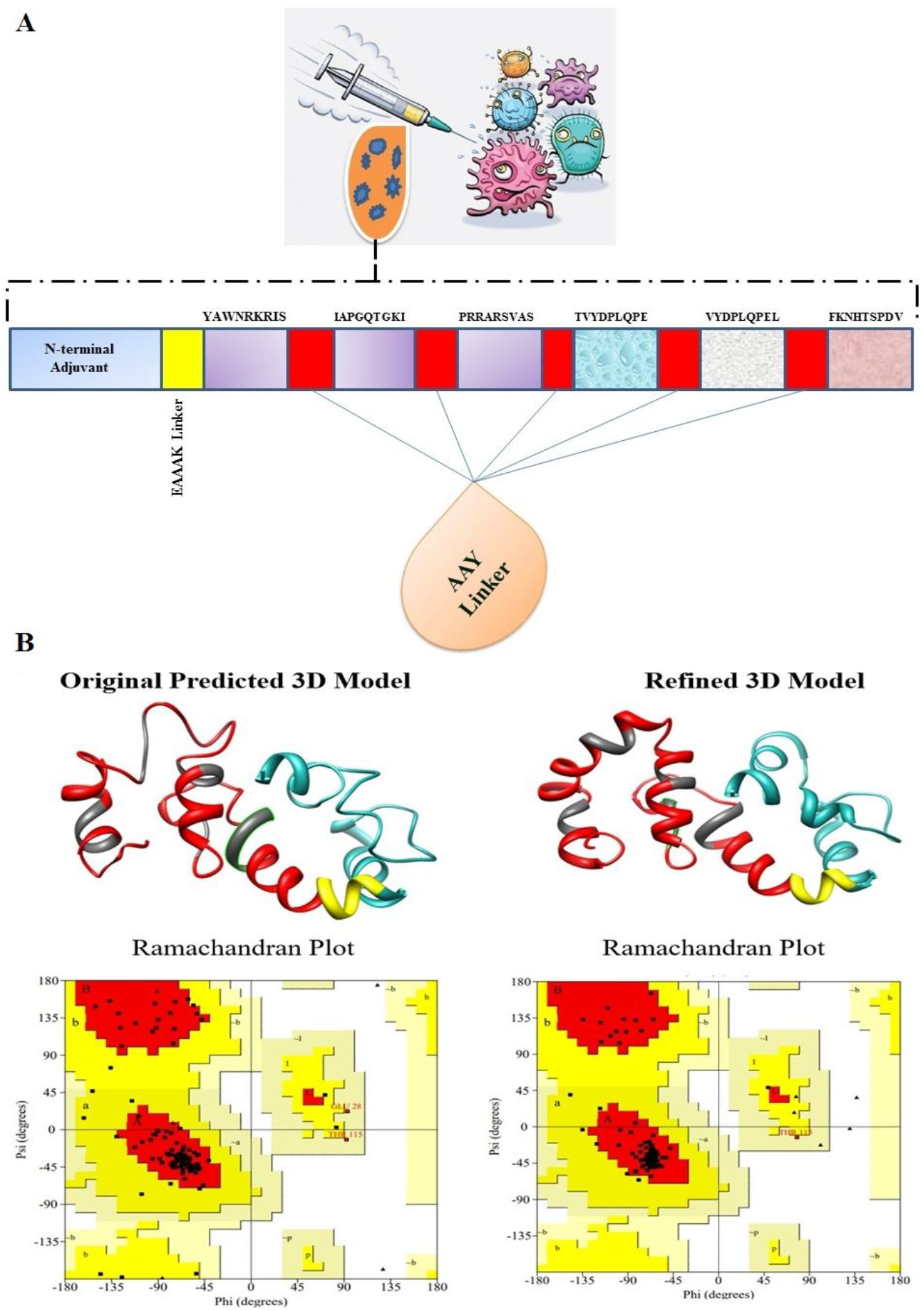
**A**. Schematic depiction of the MEPVC. **B.** The original predicted 3D MEPVC structure and refined along with respective Ramachandran plots. AAY linkers are shown in red while epitopes are in coal and yellow is for EAAAK linker. Cyan color represents the β-defensin adjuvant. In the Ramachandran plot, the torsion angles are shown by black squares dispersed across the core secondary structures (colored as red). The allowed regions can be understand by yellow, generously allowed by pale yellow and disallowed by white region. The top right, top left, bottom right and bottom left represent quadrants for left handed alpha helices, beta sheets, right handed alpha helix, and no elements, respectively.

### 3.3. MEPVC has Appropriate Biophysicochemical Properties

Physicochemical properties of the MEPVC were evaluated in order to assist experimentalists in the field to setup experiments accordingly *in vitro* and *in vivo.* The length of MEPVC is spanned across 110 amino acids and has molecular weight of 13.30 kDa. Vaccine construct with weight less than 110 kDa is generally believed to effective vaccine target because of its easier purification. Theoretical pI of MEPVC is 9.8 and aids in locating MEPVC on 2D gel. MEPVC aliphatic index is 69.08 projecting the vaccine thermostable at different temperatures. The total number of negatively charged and positively charges residues are 8 and 22, respectively. The grand average of hydropathicity (GRAVY) score computed for the MEPVC is −0.545, illustrating hydrophilic nature of the protein and is likely to interact with water molecules. The estimated half-life of MEPVC in mammalian reticulocytes, *in vitro* is 30 hours, yeast, *in vivo* is greater than 20 hours, and *Escherichia coli, in vivo* higher than 10 hours. The antigenicity of the MEPVC was cross-checked and predicted highly antigenic with value of 0.69. Total entropy of the protein is 17.0170 which is considered ideal and also the vaccine has no transmembrane helices (alpha helical transmembrane protein, 0.0474783 and beta barrel transmembrane protein, 0.0060384) hence no difficulties can be anticipated in cloning and expression analysis. The predicted solubility upon overexpression of MEPVC is 0.965751 reflecting higher solubility of MEPVC.

### 3.4. Modeling and Refinement of MEPVC

The 3D model of the MEPVC was constructed using *ab initio* 3Dpro predictor as no appropriate template was available for homology modeling and threading methods. The 3D structure of MEPVC is shown in **Fig.5B**. The structure secured 85.4% of residues in the Ramachandran favored, 12.6%, 1.9% and 0% residues in additionally allowed, generously allowed and disallowed regions, respectively. As the predicted MEPVC unit has number of loop regions that need to be modeled proper before moving forward. In total, five sets of residues: Alas7-Lys32, Ile63-Gly69, Cys73-Arg77, Thr87-Pro102, and Asn113-Val119 were loop modeled. The loop modeled structure increased the Ramachandran favored residues percentage to 92.3%, residues in allowed region reduced to 6.7%, residues of generously allowed region to 1.0% and disallowed remained to 0%. The structure was subjected to structure perturbations and relaxations to obtain a refined model. Among the generated structures (**S-Table 2**), the first model was selected as it has improved Rama favored score, lowest stable galaxy energy of 0.96, improved clash score of 23.1 and good MolProbity value. Similarly, the structure lacks poor rotamers in contrast to the original structure. The Ramachandran statistics for the refined 3D structure are in following order: Ramachandran favored residues (93.2%), additionally allowed region (5.8%), generously allowed region (1.0%) and disallowed region (0 %). The Z-score of the refined MEPVC is −4.3 and within the score range of same size protein in structure data bases (**S-Fig.1**).

### 3.5. Disulfide engineering and *in silico* cloning of the MEPVC

Further, disulfide engineering of the MEPVC was performed in order to optimize molecular interactions and confer considerable stability by attaining precise geometric conformation [89,90]. Eight pairs of residues were selected to be replaced with cysteine amino acid. These pairs are: Gln7-Ala19 (χ3 angle,+118, energy value, 4.20 kcal/mol), Cys18-Leu21 (χ3 angle,+84.35, energy value, 3.69 kcal/mol), Lys44-Ala47 (χ3 angle,+74.17, energy value,5.59 kcal/mol), Arg57-Ala61 (χ3 angle,+122.71, energy value,6.14 kcal/mol), Ala72-Ala85 (χ3 angle,-62.92, energy value,4.40 kcal/mol), Ala73-Ala82 (χ3 angle,, energy value, kcal/mol), Leu92-Glu95 (χ3 angle, −102.37, energy value, 3.86 kcal/mol), and Phe111-Pro117 (χ3 angle,-96.0, energy value,4.14 kcal/mol). These residues have either higher energy level i.e. > 2 kcal/mol and χ3 angle out of range (< −87 and > + 97) were selected on purpose to make them stable. The original and disulfide mutant MEPVC structures are shown in **Fig.6**. The primary purpose of *in silico* cloning of the MEPVC was guide molecular biologist and genetic engineers about the possible cloning sites and predicted level of expression in a specific expression system for instance here in this study we used *E. coli* K12 system. Prior to cloning, reverse translation of the MEPVC sequence was conducted to have an optimized codon usage as per *E. coli* K12 to yield its max expression. The CAI value of the improved MEPVC sequence is 1 indicating ideal expression of the vaccine [91]. The GC content whereas is 53.2 % nearly to the *E. coli* K12 and range within the optimum ranged between 30 % and 70%. The cloned MEPVC is shown in **Fig.7**.

**Fig.6.**
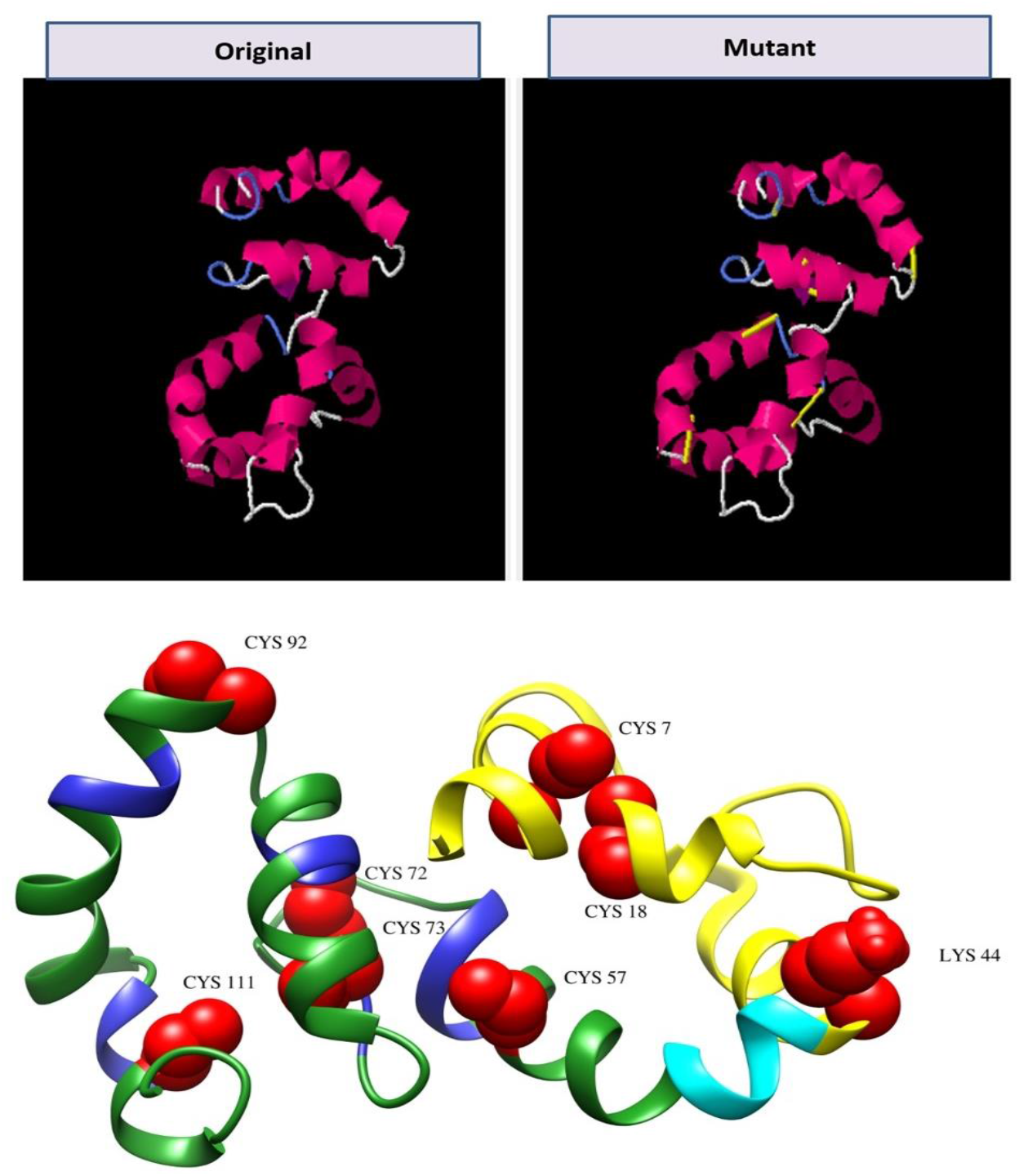
Disulfide engineering of the MEPVC (Top). The yellow bonds are the disulfide of the mutated residues to increase MEPVC stability. The red spheres in the MEPVC 3D structure (bottom) represent the mutated residues.

**Fig.7.**
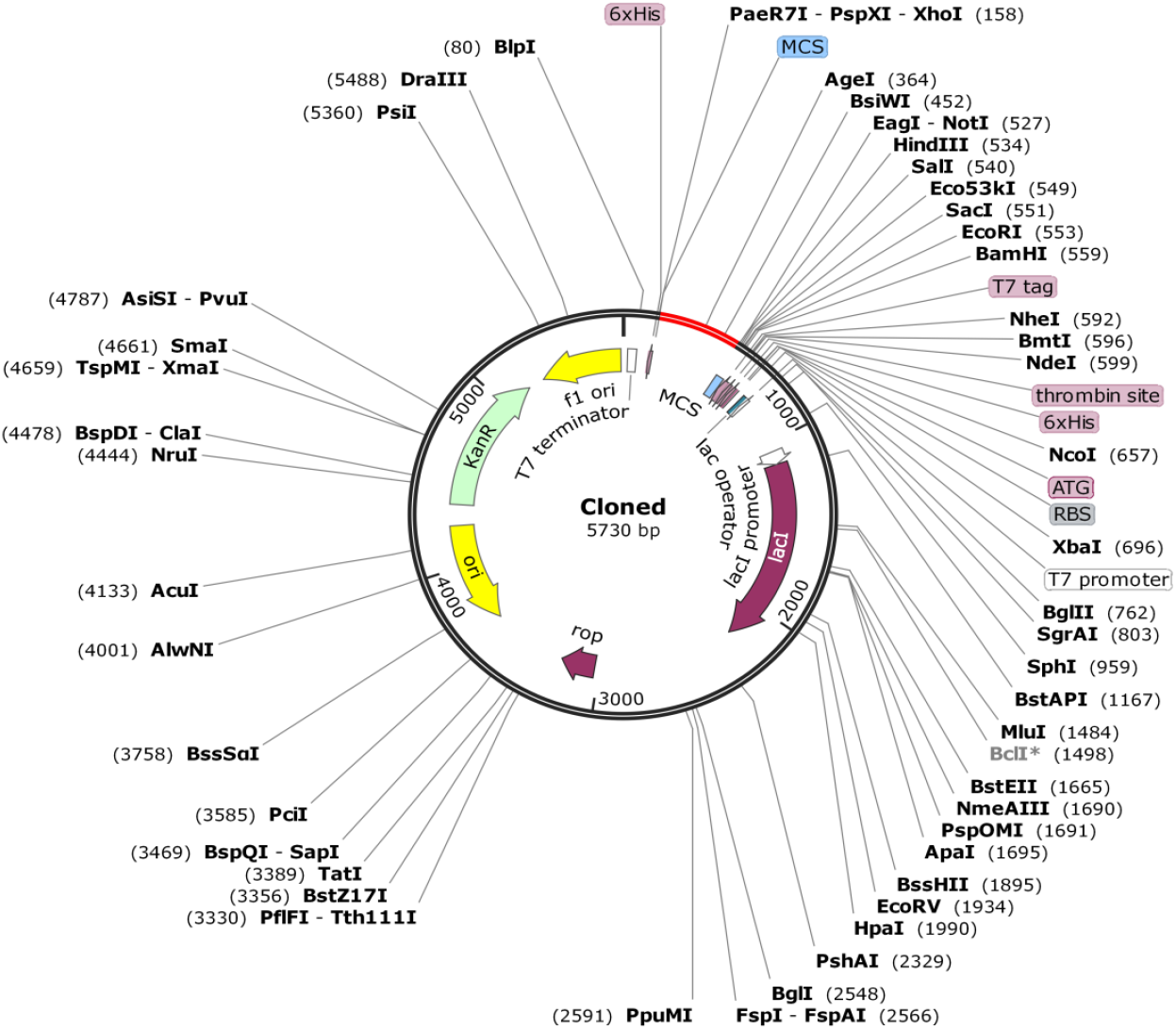
*In silico* cloning of MEPVC (shown in red) in pET28a expression vector.

### 3.6. *In silico* Immune Simulation

Both primary and secondary immune responses seem to play a significant contribution against the pathogen and may be compatible to the actual immune response. The *in silico* host immune system response to the antigen is shown in **Fig.8**. High concentration of IgG +IgG and IgM was characterized at the primary response, followed by IgM, IgG1+ IgG2 and IgG1 at both primary and secondary stages with concomitant of antigen reduction. Additionally, robust response of interleukins and cytokines were observed. All this suggest the efficient immune response and clearance of the pathogen upon subsequent encounters. Elevated B cell population including memory cells and different isotypes in response to the antigen, points to the long lasting formation of memory and isotype switching. The T helper cell population additionally with the cytotoxic T cell and their respective memory development are in strong agreement of strong response to the antigen.

**Fig.8.**
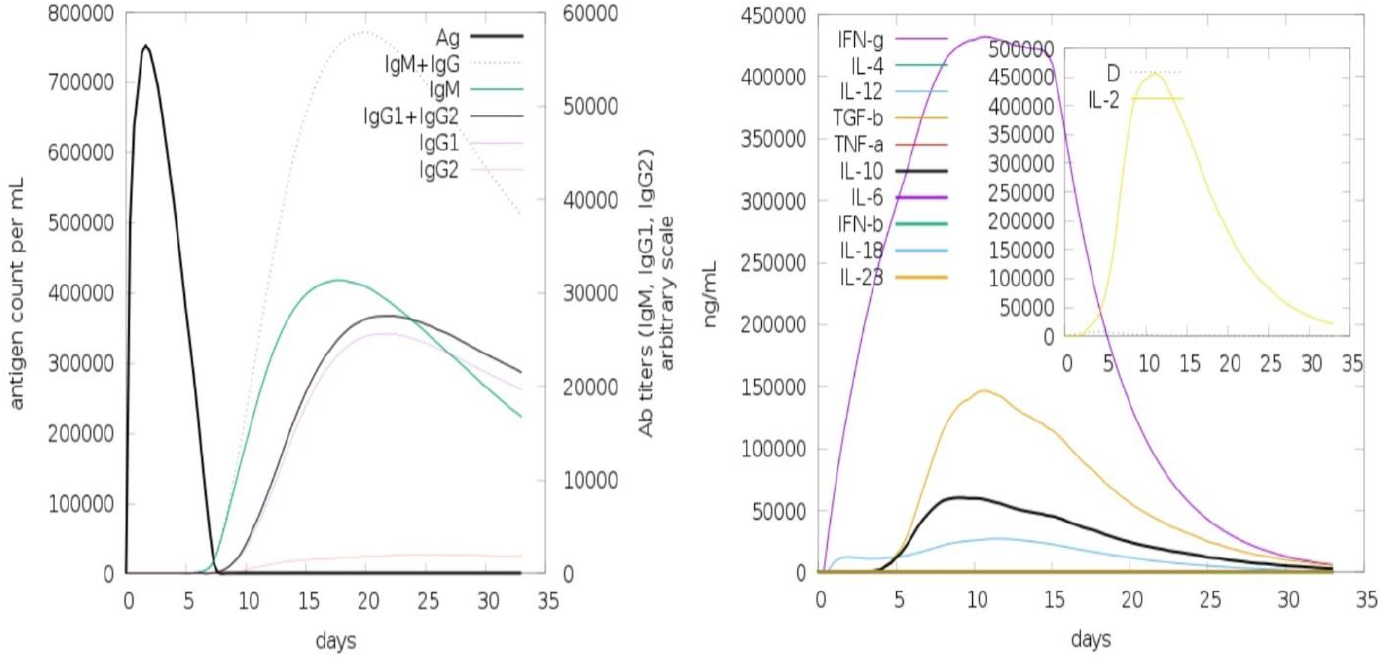
Computational immune simulation of the host immune system response to the MEPVC. The antibodies are shown in left and cytokines and interleukins in right.

### 3.7. Interaction of MEPVC and TLR3 Promotes Formation of Strong Complex

Bioinformatic modelling driven molecular docking of the desingned MEPVC to one representative innate immune response receptor TLR3 was carried out in order to decode MEPVC potential of binding to the innate immune receptors. This was fundamental to understand as TLR3 is significant in recognition of virus associated molecular patterns and of activaiton of type I interferons and NF-kappa B. The docking assesment predicted top 20 complexes sorted mainlny on scoring functions along with interacting molecules area size, desolvation energy, and complexes actual rigid transformation. Following, the complexes were subjected to FireDock web server for refinement assay. This ease in discarding flexibility errors of the docking procedure and provide a deep refinement of the predictions thus limiting the chances of false positive docking calculations. According to the global energy, solution 8 was considered as a best complex with net global energy of −20.78 kJ/mol (**Table 2**). This energy is the output of −16.88 kJ/mol attractive van der Waals (vdW), 3.81 kJ/mol repulsive vdW, 8.14 kJ/mol atomic contact energy (ACE), and −0.93 kJ/mol hydrogen bond energy. The docked conformation and chemical interacting residues of the MEPVC with TLR3 is shown in **Fig.9**. Visual inspection of the complex leads to observation of deep binding of the MEPVC at the center of TLR3 and favor rigorously rigoursly hydrogen and weak van dar Waals interactions with various residues of TLR3. Within 3 Å, the MEPVC was noticed to formed interactions with His39,Val55,Asn57,Asp81,Phe84,Val103,Asn105,Gln107,His108,Thr126,Glu127,Ser132,Hi s129,Thr151,His156,Gln174,GLu175,Lys200,Lys201,Glu203,Asn229,Ser256,Asp280,Ser282,Tyr283,Tyr302,Phe304,Tyr307,Lys330,Tyr383,Tyr326,Asn328,His359,Asn361, and Glu363.

**Fig.9.**
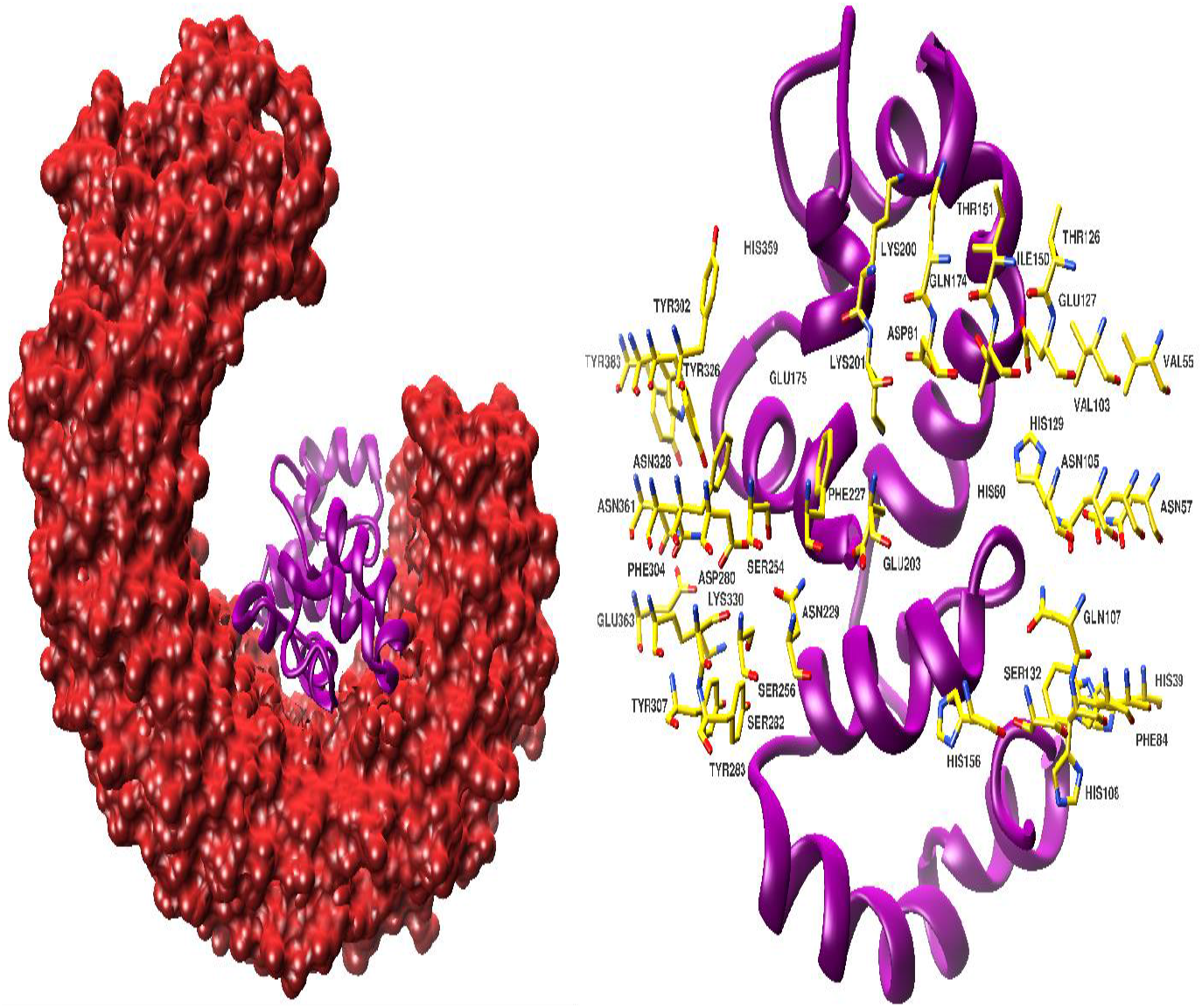
**A**. MEPVC (shown in dark magenta) conformation with respect to TLR3 receptor (shown in firebrick surface). **B**. TLR3 interacting residues (shown in yellow sticks) are involved in hydrophobic and hydrophilic interactions with MEPVC.

**Table 2.**
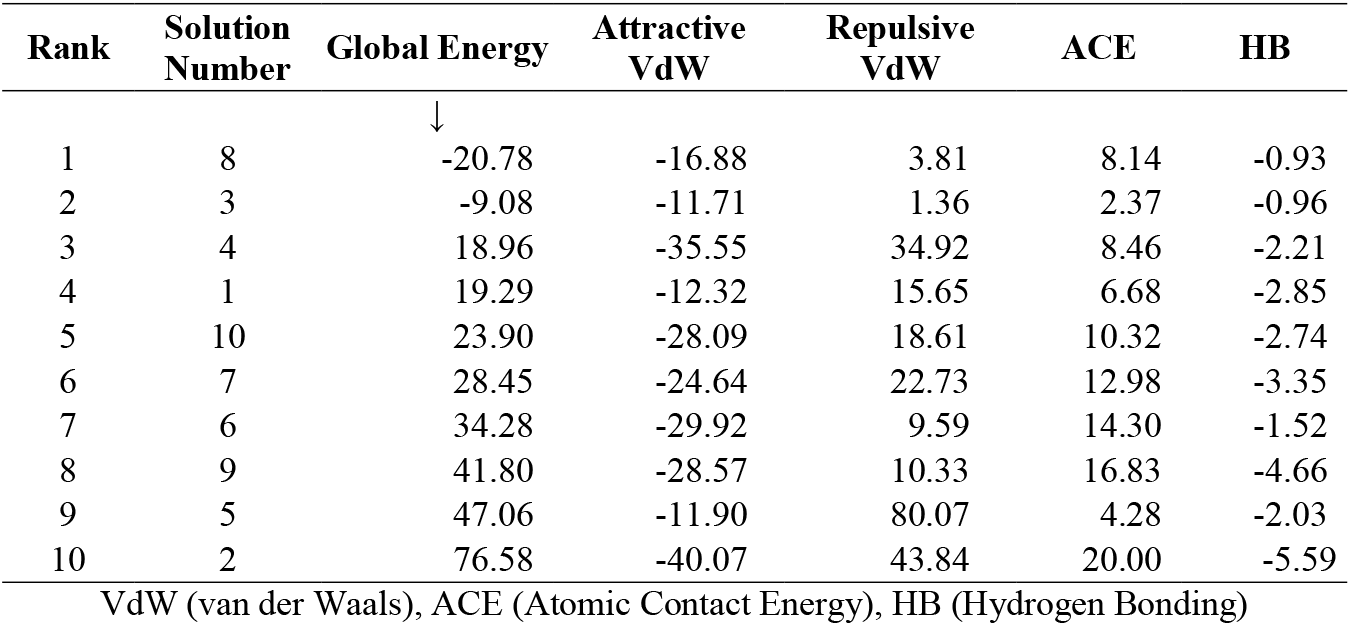
Refined PatchDock complexes as an outcome of FIreDock assay.

| Rank | Solution Number | Global Energy | Attractive VdW | Repulsive VdW | ACE | HB |
| --- | --- | --- | --- | --- | --- | --- |
|  |  | ↓ |  |  |  |  |
| 1 | 8 | -20.78 | -16.88 | 3.81 | 8.14 | -0.93 |
| 2 | 3 | -9.08 | -11.71 | 1.36 | 2.37 | -0.96 |
| 3 | 4 | 18.96 | -35.55 | 34.92 | 8.46 | -2.21 |
| 4 | 1 | 19.29 | -12.32 | 15.65 | 6.68 | -2.85 |
| 5 | 10 | 23.90 | -28.09 | 18.61 | 10.32 | -2.74 |
| 6 | 7 | 28.45 | -24.64 | 22.73 | 12.98 | -3.35 |
| 7 | 6 | 34.28 | -29.92 | 9.59 | 14.30 | -1.52 |
| 8 | 9 | 41.80 | -28.57 | 10.33 | 16.83 | -4.66 |
| 9 | 5 | 47.06 | -11.90 | 80.07 | 4.28 | -2.03 |
| 10 | 2 | 76.58 | -40.07 | 43.84 | 20.00 | -5.59 |

VdW (van der Waals), ACE (Atomic Contact Energy), HB (Hydrogen Bonding)

### 3.8 MD Simulations Supported Docked Complex Stability and Equilibrium

The stability of MEPVC with TLR3 was further investigated through MD simulations. The trajectories of MD simulations were used in vital statistical analysis to decode backbone stability and residual flexibility. Root mean square deviation (RMSD) [92,93] was performed first that compute average distance of backbone carbon alpha atoms of superimposed frames (**Fig.10A**). The average RMSD for the system is 3.23 Å with max of 4.4 Å at 24-ns. An initial sudden change in RMSD can be seen up to 2.7-ns that may be due to adjustments adopted by the complex when exposed to dynamics forces and milieu. The second minor RMSD shift can be noticed between 22-ns to 26-ns. Afterward, the system is quite stable with not global and local conformational changes specified. Next, root mean square fluctuations (RMSF) [94] was applied on the system trajectories (**Fig.10B**). RMSF is the average residual mobility of complex residues from its mean position. Mean RMSF for the MEPVC-TLR3 complex calculated is 1.60 Å with max of 8.6 Å pointed at the N-terminal of the MEPVC. Most of the interacting residues of the MEPVC with TLR3 are subject to minor fluctuations, a fact in analogy to complex high stability. The thermal residual deviation was assessed afterward by beta-factor (β-factor) [95], the outcomes of which is strongly correlated to the RMSF and hence further affirming system stability (**Fig.10C**). The average β-factor of the system analyzed is 88.64 Å^2^ with max of 1956.23 Å^2^. Lastly, we evaluated the compactness of the system by means of radius of gyration (Rg) [96] analysis (**Fig.10D**). High Rg and low R*g* illustrate the magnitude of system compactness and system less tight packing. It further tell us the whether the system of interest in order or not. Highly compact system is an indication of system stability and vice versa. The mean Rg for our system is 55.8411 Å with max score of 74.884 reflecting higher ordered and compact nature of the system.

**Fig.10.**
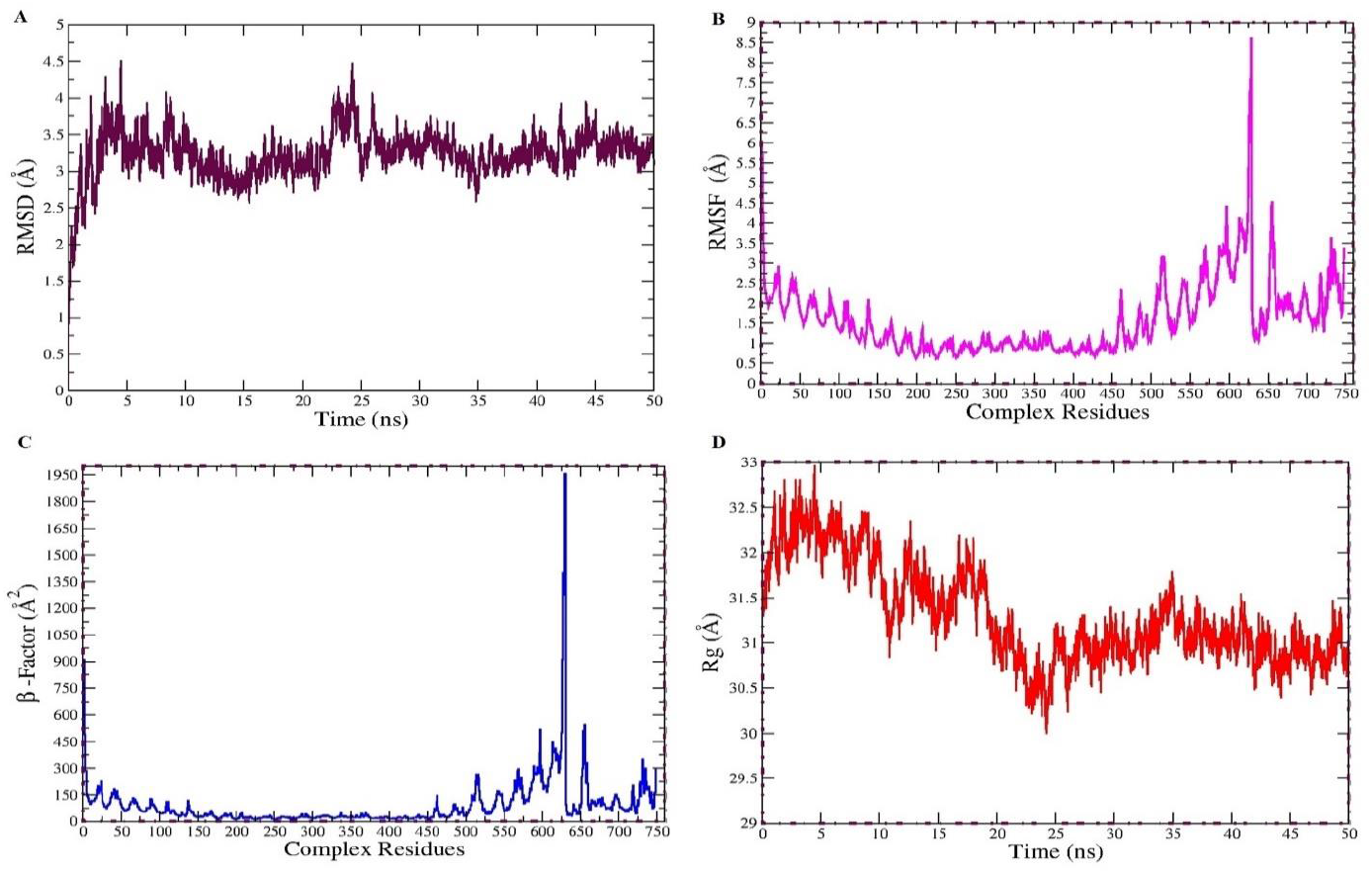
MD simulations based analysis of RMSD (A), RMSF (B), β-factor (C), and Rg (D).

### 3.9. Hydrogen bonds Analysis

Hydrogen bonds are dipole-dipole attractive forces and formed when a hydrogen atom bounded to a highly electronegative atom such as F, N, and O is attracted by another electronegative atom [97]. The strength of a hydrogen bond vary from 4 kJ to 50 kJ per mole. Hydrogen bonds are deemed vital in molecular recognition and provide rigidity in achieving stable conformation [98]. The frequency of hydrogen bonds in each frame of the MD simulation trajectories can be visualized in **Fig.11A**. These hydrogen bonds are extracted by mean of VMD HBonds plugin and are 104 in number as tabulated in **Table 3**. The cut-off distance set is 3.0 Å and cut-off angle 20 degrees. Each residue pair may for one, two or more each of which is counted separately. The min, mean and max number of hydrogen bonds between MEPVC and TLR3 are 1, 5, and 12, respectively.

**Fig.11.**
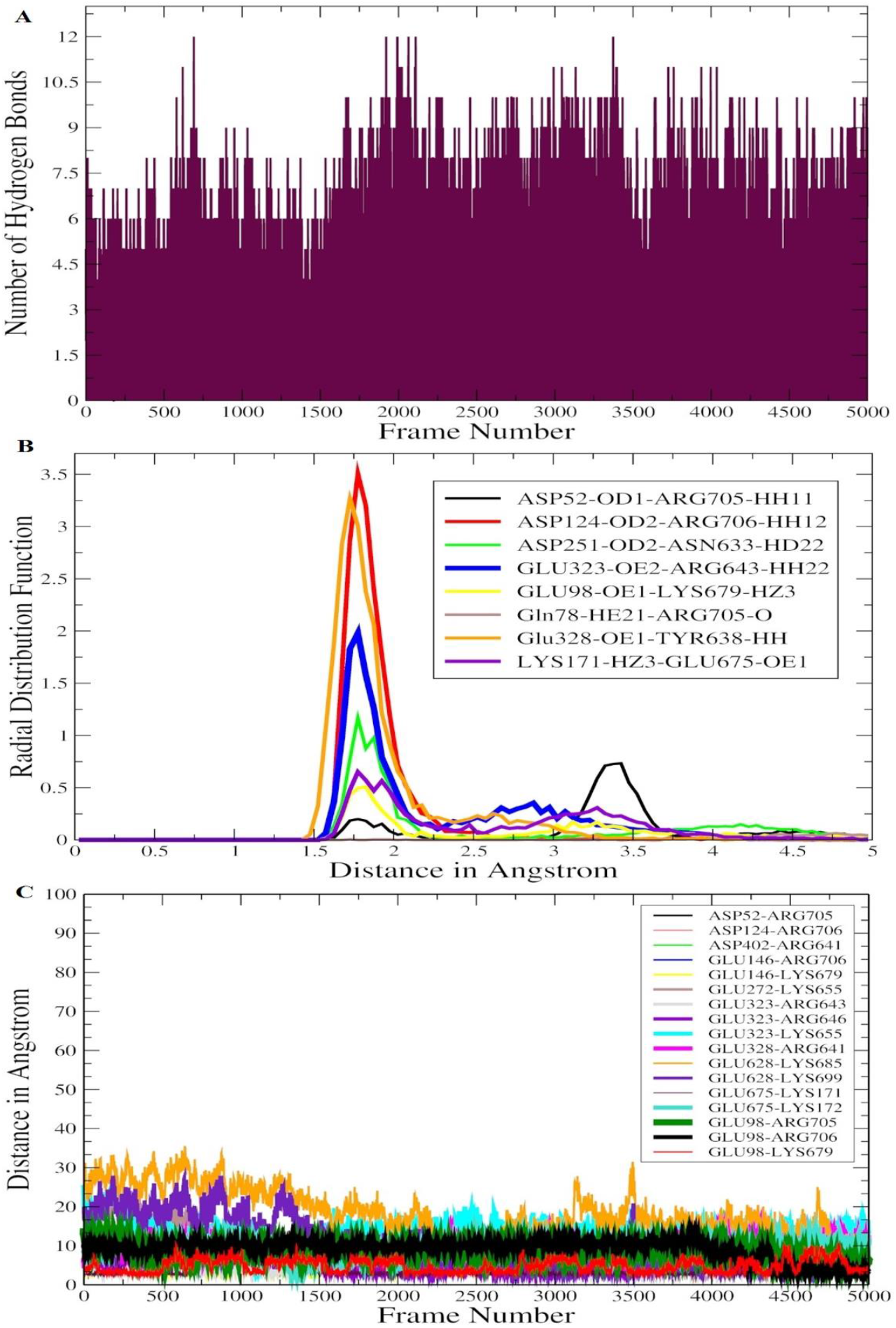
Number of hydrogen bonds in each frame of MD simulation trajectories (A), RDF plots for hydrogen bonds in TLR3-MEPVC critical in interaction and stability (B), and salt bridges formed between TLR3 and MEPVC (C) during simulation time.

**Table 3.**
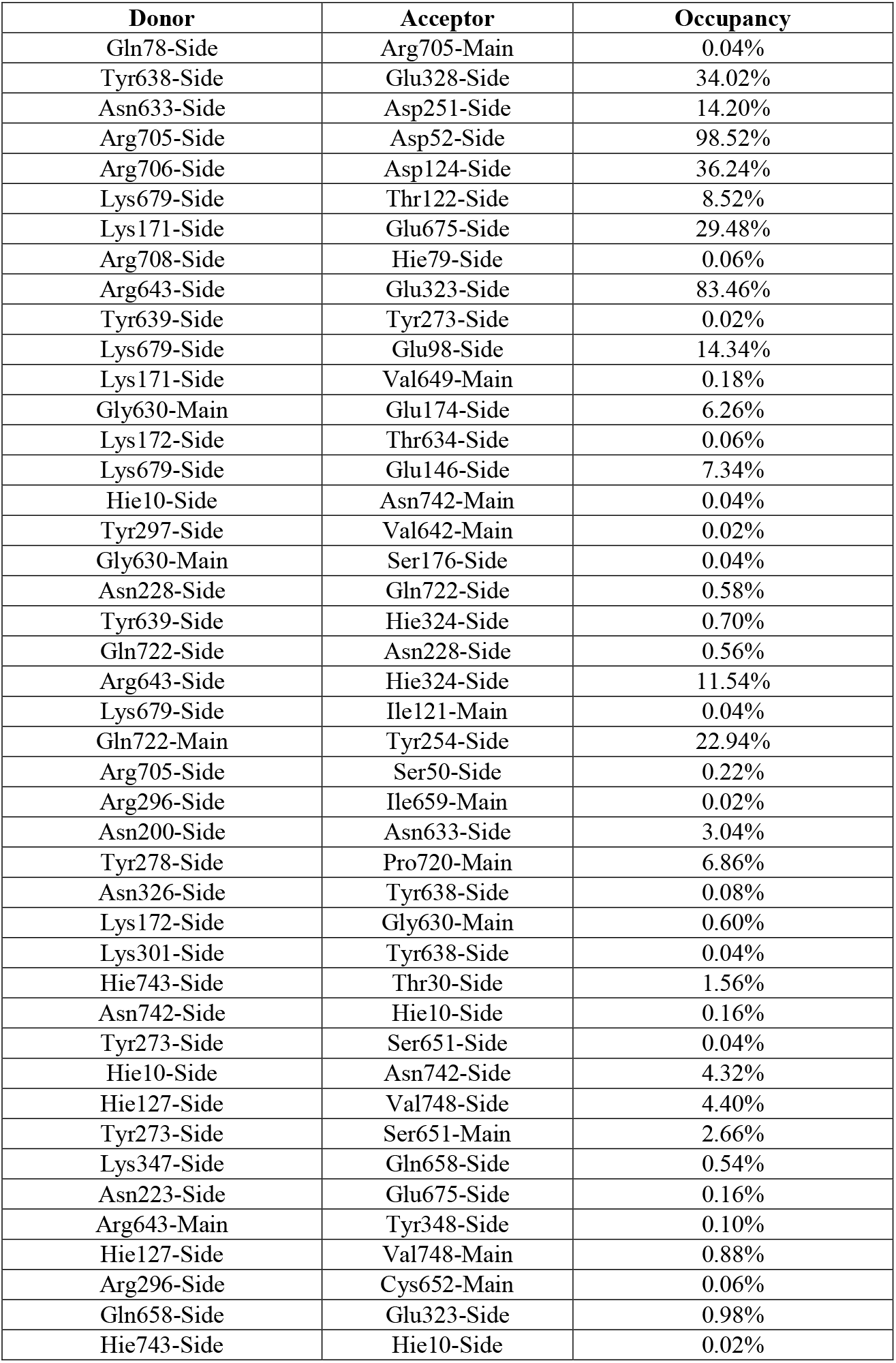
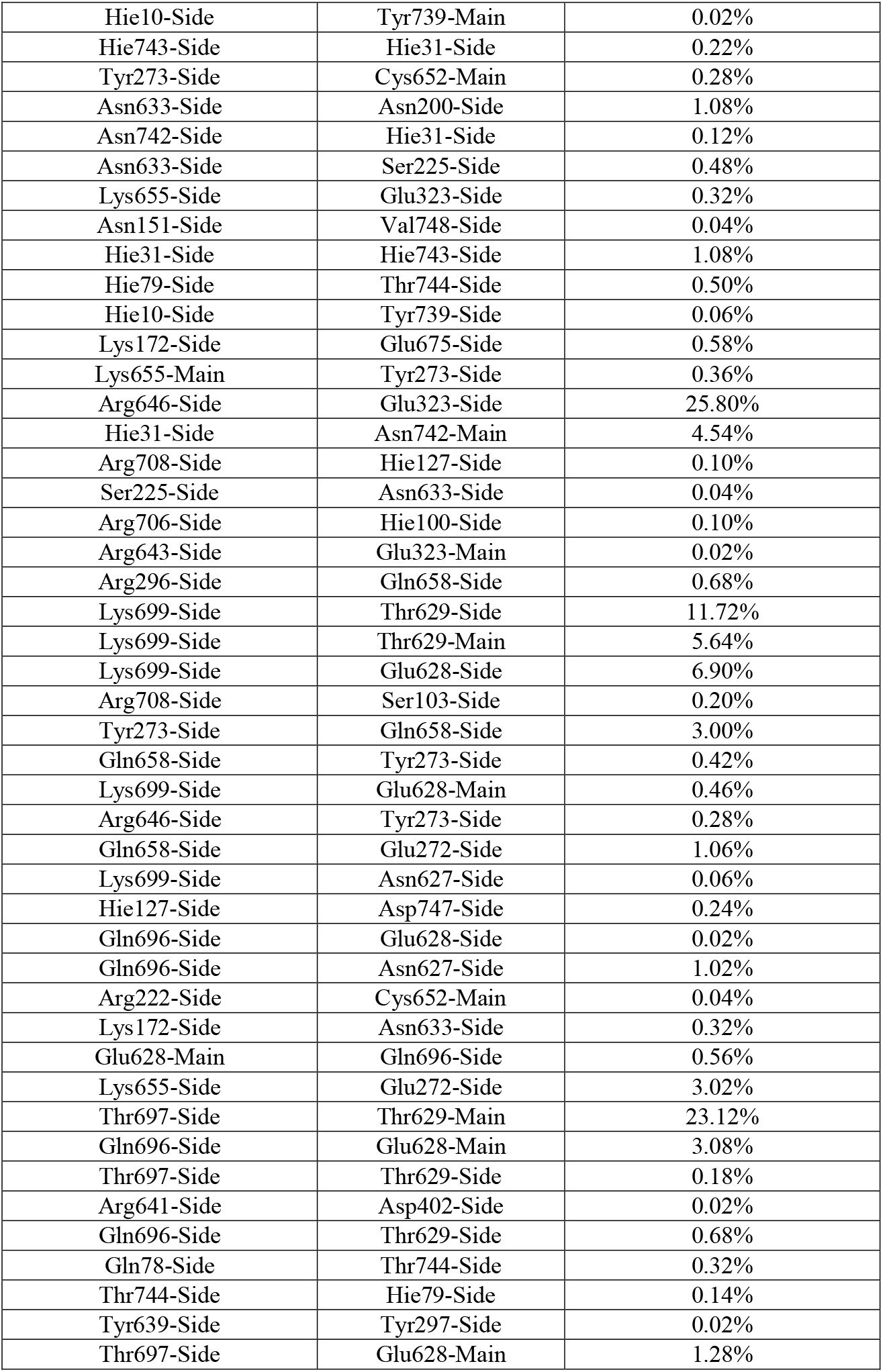
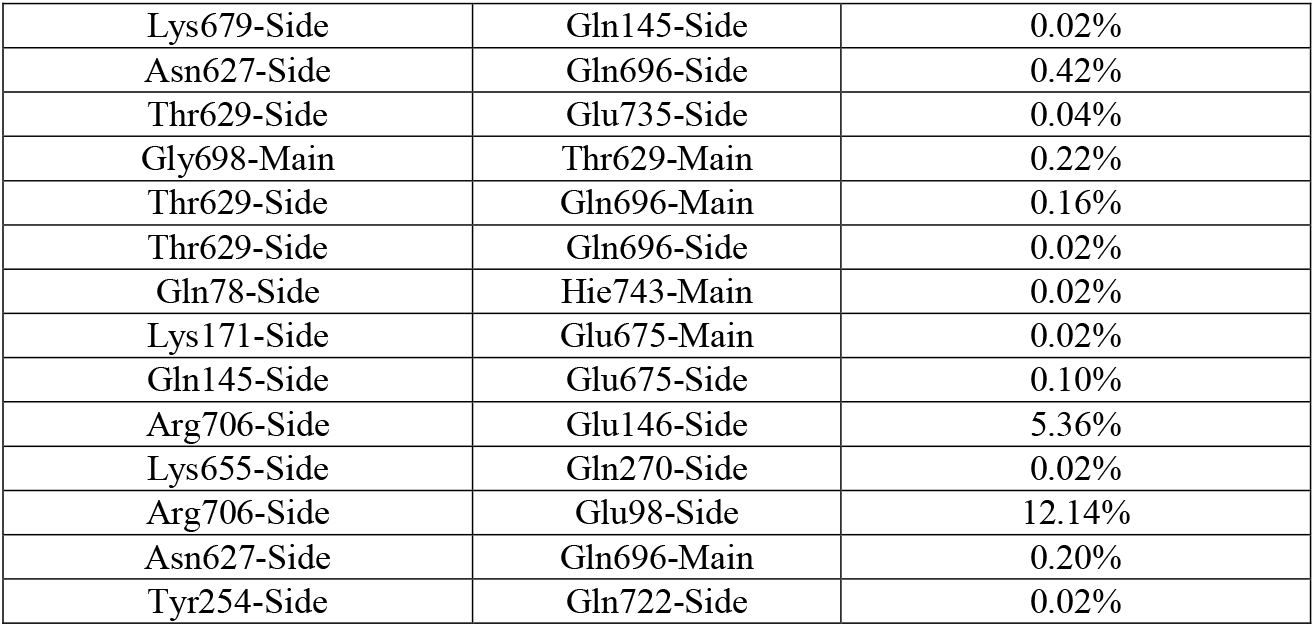
Hydrogen bonds between TLR3 and MEPVC formed during MD simulations.

### 3.10. Radial Distribution Function and Inter Molecular Interactions

The distribution and bonding pattern of intermolecular interactions of the MEPVC residues atom(s) with respect to the TLR3 were studied through radial distribution function (RDF) (Abbasi et al., 2016; Donohue, 1954; Kouetcha et al., 2017). RDF mainly describes distance ‘r’ between two entities and is represented by g(r). The factor ‘r’ is extracted from simulation trajectories and range from o to ∞ [75]. The hydrogen bonds predicted by VMD were utilized in RDF that shown only 8 interactions between MEPVC and TLR3 with good affinity for each other. In these interactions, TLR3 residues (atoms) are: Asp52:OD1, Gln78:HE21, Glu98:OE1, Asp124:OD2, Lys171:HZ3, Asp251:OD2, Glu323:OE2, and Glu328:OE1 are found to have strong radii distribution to their counterpart MEPVC residues (atoms): Arg705:HH11, Arg705: O, Lys679:HZ3, Arg706:HH12, Glu675:OE1, Asn633:HD22, Arg643:HH22, and Tyr638: HH, respectively. The RDF plots for the above said interactions are illustrated in **Fig.11B**. The interaction between Asp124-OD2 and Arg706-HH12 has a refined distribution pattern and highest density distribution among all. The max g(r) value for this interaction is 3.51 observed at distance range of 1 Å. This is followed by Glu328-OE1-Tyr638-HH with max g(r) value of 3.26 mostly interaction at distance range of 0.6 Å. The Glu323-OE2-Arg643-HH22 is also much refined having g(r) value of 1.98 and mostly interacts within distance range of 0.6 Å. The remaining interactions density distribution is not confined and vary considerably but important from MEPVC and TLR3 interaction point of view.

### 3.11 Salt Bridges and TLR3-MEPVC Stability

Salt bridges are non-covalent in nature and the outcome of interactions between two ionized states [101]. These interactions comprised two parts: an electrostatic interaction and a hydrogen bond. In salt bridges, lysine or arginine typically behave as base where glutamine or aspartate as acid and the bridge is created when carboxylic acid group allows a proton migration to guanidine and amine group in arginine. Salt bridges are the strongest among all non-covalent interactions and contribute to a major extent in biomolecular stability [102–104]. In total, 17 salt bridges were identified between TLR3 and MEPVC within the cut-off distance of 3.2 Å as can be depicted from **Fig.11C**. The higher numbers of salt bridges were recorded for TLR3-Glu628 and MEPVC-Lys685. The mean number of salt bridges for this interaction is 18, max, 35 and min, 3. The count for other salt bridges from TLR3 to MEPVC is in following order: Asp124-Arg706 (mean, 3, max,7 and min,3), Glu98-Lys679 (mean, 5, max,10 and min,4), Asp402-Arg641 (mean, 12, max, 18 and min, 4), Glu146-Arg706 (mean,7 max,11 and min,3), Glu146-Lys679 (mean,5 max,13 and min,2), Glu272-Lys655 (mean,9 max,22 and min,2), Glu323-Arg643 (mean,4 max,8 and min,3), Glu323-Arg646 (mean,5 max,10 and min,3), Glu323-Lys655 (mean,13 max,25 and min,3), Glu328-Arg641 (mean,10 max,17 and min,4), Glu628-Lys699 (mean,9 max,28 and min,2), Glu675-Lys171 (mean,4 max,13 and min,2), Glu675-Lys172 (mean,10 max,16 and min,2), Glu98-Arg705 (mean,9 max,14 and min, 5), Glu98-Arg706 (mean, 9 max,14 and min,3), and Glu98-Lys679 (mean, 5 max,11 and min,2).

### 3.12. Density Distribution and Local Structure Movements

The vital hydrogen bond interactions involved between TLR3 receptor and MEPVC shortlisted by VMD were subjected to a novel AFD analysis to elucidate 3D movements of MEPVC atoms with respect to a reference TLR3 residues atom in simulation time. To this objective, interactions mentioned in the RDF were used in AFD. Preliminary investigation suggested that only three interactions: TLR3-Asp52-MEPVC-Arg705, TLR3-Glu328-MEPVC-Tyr638, and TLR3-Glu323-MEPVC-Arg643 are mainly represented frequently and found in most of the simulation frames. The TLR3-Asp52-MEPVC-Arg705 is uncovered in 4997 frames, TLR3-Glu328-MEPVC-Tyr638 in 4988, and TLR3-Glu323-MEPVC-Arg643 in 4985 making these interactions ideal for interpreting density distribution of the interactions on XYZ planes and also appropriate for gaining ideas about conformational changes of the interacting atoms with respect to each other. As the local structure movements and rotations are responsible for functional shifts, their understanding in our system is important to be unveiled. For TLR3-Asp52-MEPVC-Arg705 (**Fig.12**), the density distribution is not uniform, dispersed and behave flexibility in affinity on all three axis for the receptor atom. Parallel, the strength of interaction is also observed affected due to these minor structural movements of the MEPVC residue atom. Though, the mentioned interaction depicts MEPVC is still within the vicinity of the TLR3 reference residue and enjoys this interaction flexibility with the said MEPVC residue during simulation. TLR3-Glu328-MEPVC-Tyr638 interaction (**Fig.13**) has less distribution area and has much higher intensity illustrating strong affinity of the interacting atoms for each other. It also gives an idea of the lesser movements of the atoms with respect to each other, an indication of a correct system conformation. The distribution area TLR3-Glu323-MEPVC-Arg643 is much dispersed though high intensity of the interaction can be seen in close vicinity (**Fig.14**).

**Fig.12.**
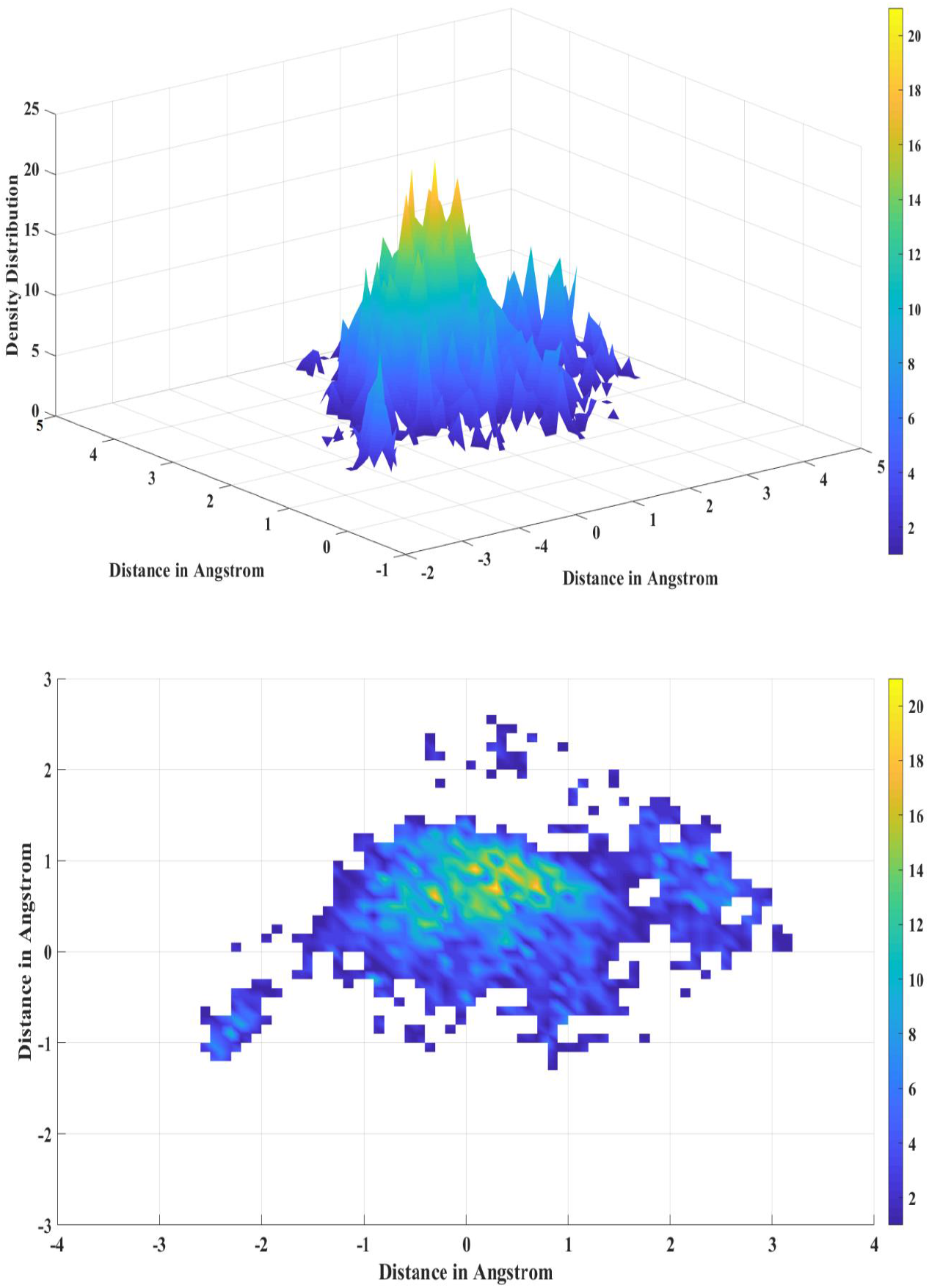
AFD for TLR3-Asp52-MEPVC-Arg705 interaction.

**Fig.13.**
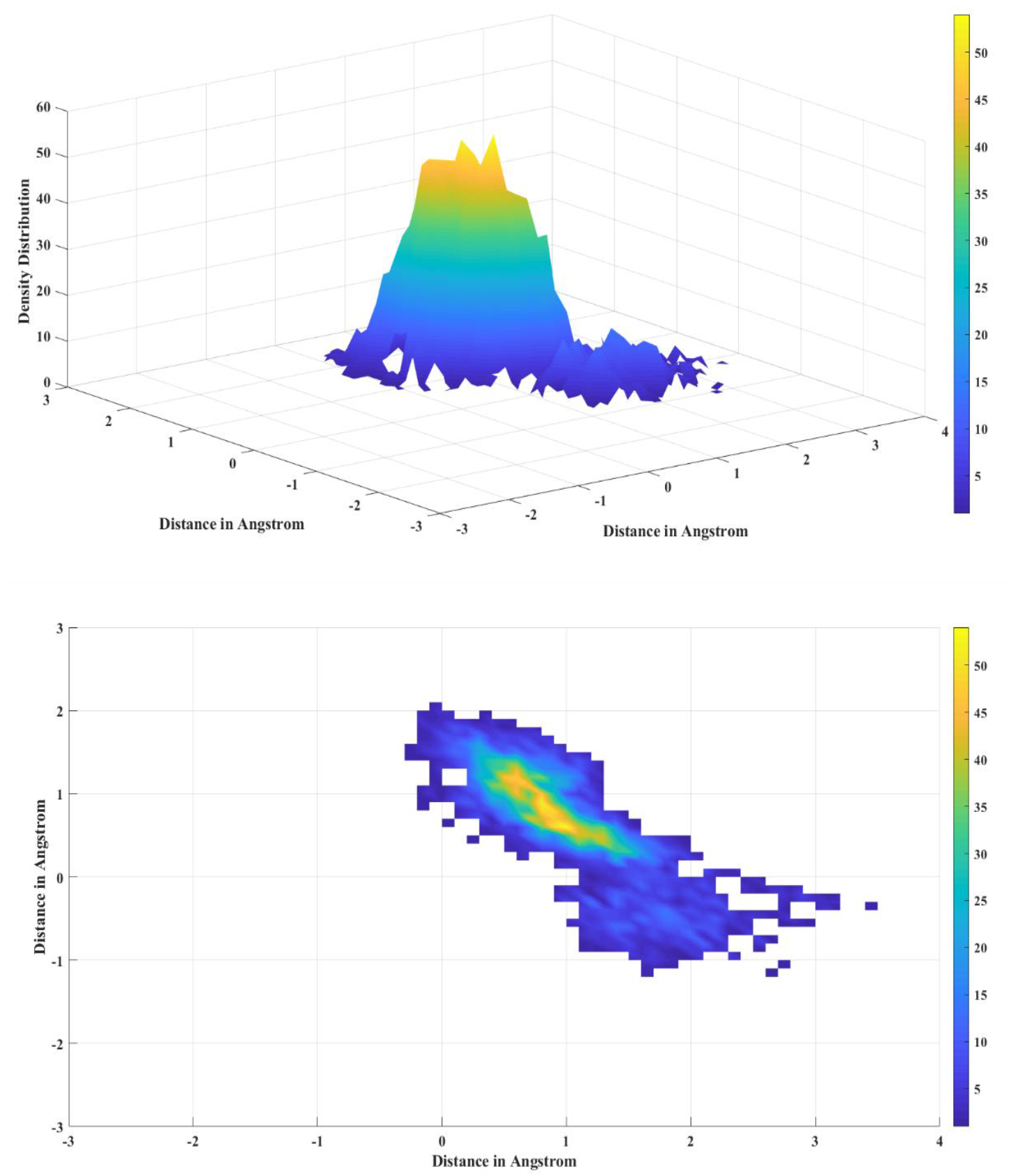
AFD plot for TLR3-Glu328-MEPVC-Tyr638.

**Fig.14.**
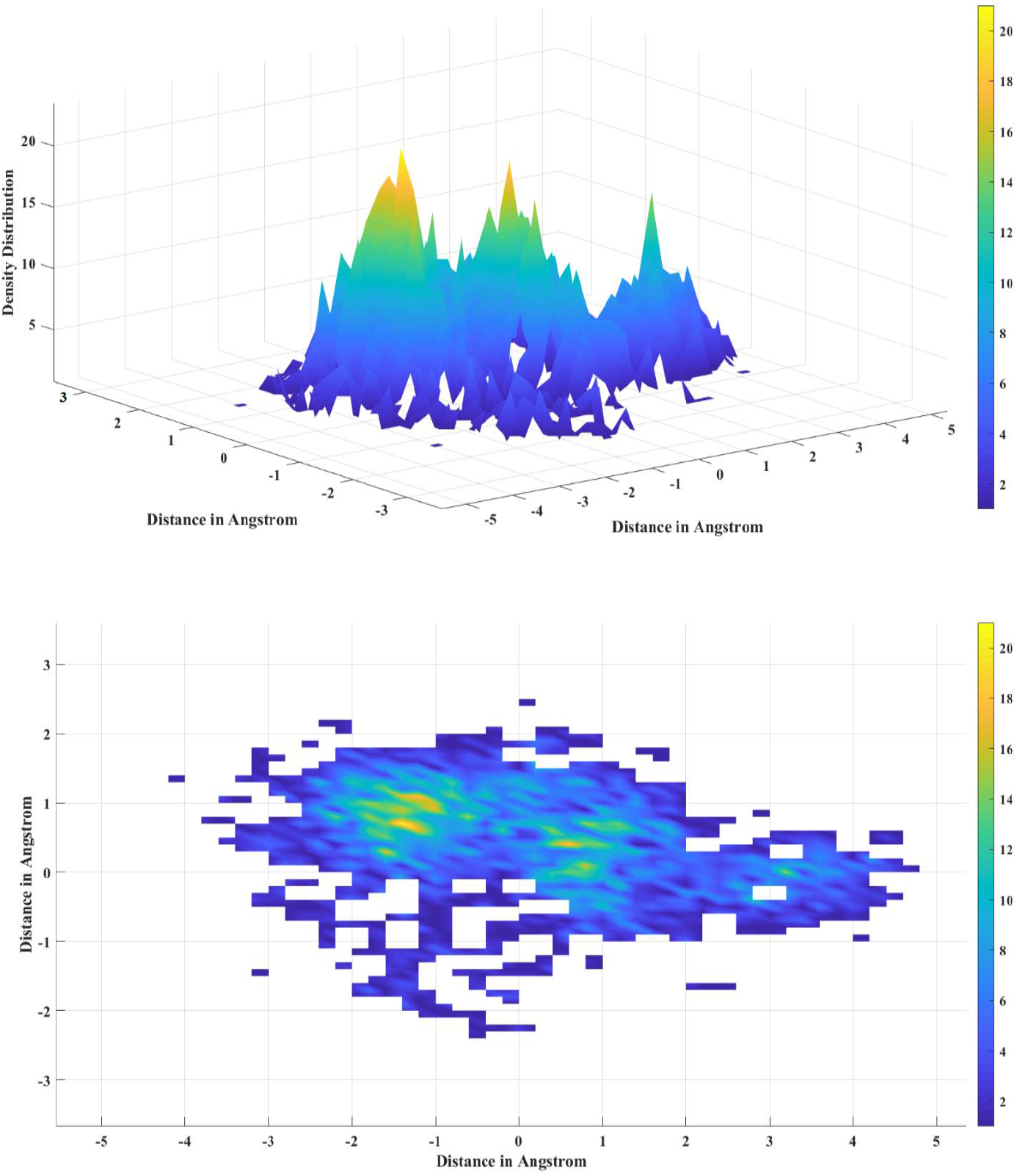
AFD plot for TLR3-Glu323-MEPVC-Arg643.

### 3.13. MEPVC-TLR3 Complex Revealed High Binding Free Energies

The net free energy of binding (ΔTOTAL) in both GB and PB models are revealed favorable MEPVC-TLR3 complex in pure water. The net GB and PB energy for the MEPVC-TLR3 complex is −53.81 kcal/mol and −89.02 kcal/mol, respectively. To this energy, high contribution was noticed from gas phase energy (ΔG gas) compared to highly insignificant contributions from solvation energy (ΔG solv). In GB model, the ΔG gas energy for the system is −1889.76 kcal/mol whereas in PB model this energy is −1889.76 kcal/mol. The ΔG solv energy in case of GB is 1835.95 kcal/mol while in case of PB it is 1800.74 kcal/mol. The electrostatic contribution to the system estimated by MM force field in both methods is highly favorable to the net energy (i.e. −1752.51 kcal/mol). Likewise, the van der Waals contribution form MM is also reported significant in system stability (−137.25 kcal/mol). The electrostatic energy contribution (EGB and EPB) to the ΔG solv is found prime parameter leading to non-favorable contribution of ΔG solv energy in both techniques. The surface area energy (ESURF) computed in GB method is −20.22 kca/mol. In PB, ENPOLAR and EDISPER are the repulsive and attractive free energy and is −17.44 kcal/mol and 0 kcal/mol, respectively. Individual binding free energy for MEPVC-TLR3 complex, TLR3 receptor, MEPVC and net energy is tabulated in **Table 4**.

**Table 4.** Binding free energies for MEPVC-TLR3 system.

| GB |  |  |  | PB |  |  |  |
| --- | --- | --- | --- | --- | --- | --- | --- |
| MEPVC-TLR3 Complex |  |  |  | MEPVC-TLR3 Complex |  |  |  |
| Energy Component | Average | Std. Dev. | Std. Err. of Mean | Energy Component | Average | Std. Dev. | Std. Err. of Mean |
| VDWAALS | -6191.82 | 42.04 | 4.2 | VDWAALS | -6191.82 | 42.04 | 4.2 |
| EEL | -51977.55 | 163.19 | 16.31 | EEL | -51977.55 | 163.19 | 16.31 |
| EGB | -9135.8 | 129.32 | 12.93 | EPB | -8914.84 | 123.4 | 12.34 |
| ESURF | 236.0035 | 2.49 | 0.24 | ENPOLAR | 162.86 | 1.12 | 0.11 |
| G gas | -58169.38 | 155 | 15.5 | G gas | -58169.38 | 155 | 15.5 |
| G solv | -8899.79 | 128.24 | 12.82 | G solv | -8751.97 | 123.14 | 12.31 |
| TOTAL | -67069.18 | 66.36 | 6.63 | TOTAL | -66921.36 | 69.42 | 6.94 |
| TLR3 Receptor |  |  |  | TLR3 Receptor |  |  |  |
| Energy Component | Average | Std. Dev. | Std. Err. of Mean | Energy Component | Average | Std. Dev. | Std. Err. of Mean |
| VDWAALS | -5255.53 | 38.07 | 3.8 | VDWAALS | -5255.53 | 38.07 | 3.807 |
| EEL | -44981.18 | 160.75 | 16.07 | EEL | -44981.1 | 160.75 | 16.07 |
| EGB | -7204.15 | 129.25 | 12.92 | EPB | -6992.97 | 121.61 | 12.16 |
| ESURF | 199.02 | 2.05 | 0.2 | ENPOLAR | 140.11 | 0.93 | 0.09 |
| G gas | -50236.72 | 147.41 | 14.74 | G gas | -50236.72 | 147.41 | 14.74 |
| G solv | -7018 | 128.74 | 12.87 | G solv | -6852.86 | 121.47 | 12.14 |
| TOTAL | -57241.85 | 54.49 | 5.44 | TOTAL | -57089.58 | 60.3 | 6.03 |
| MEPVC |  |  |  | MEPVC |  |  |  |
| Energy Component | Average | Std. Dev. | Std. Err. of Mean | Energy Component | Average | Std. Dev. | Std. Err. of Mean |
| VDWAALS | -799.03 | 15.45 | 1.54 | VDWAALS | -799.03 | 15.45 | 1.54 |
| EEL | -5243.85 | 56.53 | 5.65 | EEL | -5243.85 | 56.53 | 5.65 |
| EGB | -3787.82 | 47.6 | 4.76 | EPB | -3740.06 | 45.41 | 4.54 |
| ESURF | 57.2 | 1.08 | 0.1 | ENPOLAR | 40.2 | 0.64 | 0.06 |
| G gas | -6042.89 | 56.11 | 5.61 | G gas | -6042.89 | 56.11 | 5.61 |
| G solv | -3730.62 | 47.43 | 4.74 | G solv | -3699.86 | 45.23 | 4.52 |
| TOTAL | -9773.51 | 31.87 | 3.18 | TOTAL | -9742.75 | 33.63 | 3.36 |
| Differences (MEPVC-TLR3 - TLR3- Vaccine MEPVC) |  |  |  | Differences (MEPVC-TLR3 - TLR3- Vaccine MEPVC) |  |  |  |
| Energy Component | Average | Std. Dev. | Std. Err. of Mean | Energy Component | Average | Std. Dev. | Std. Err. of Mean |
| VDWAALS | -137.25 | 7.9 | 0.79 | VDWAALS | -137.25 | 7.9 | 0.79 |
| EEL | -1752.51 | 48.84 | 4.88 | EEL | -1752.51 | 48.84 | 4.88 |
| EGB | 1856.18 | 45.486 | 4.54 | EPB | 1818.19 | 45.33 | 4.53 |
| ESURF | -20.22 | 1.29 | 0.12 | ENPOLAR | -17.44 | 0.51 | 0.05 |
| - | - | - | - | EDISPER | 0 | 0 | 0 |
| $\Delta G_{\text{gas}}$ | -1889.76 | 48.37 | 4.83 | $\Delta G_{\text{gas}}$ | -1889.76 | 48.37 | 4.83 |
| $\Delta G_{\text{solv}}$ | 1835.95 | 45.19 | 4.51 | $\Delta G_{\text{solv}}$ | 1800.74 | 45.2 | 4.52 |
| $\Delta \text{TOTAL}$ | -53.81 | 8.1955 | 0.8195 | $\Delta \text{TOTAL}$ | -89.02 | 11.55 | 1.15 |

### 3.14. Net Energy Decomposition Discovered Hot-spot Residues

The net free energy of the simulated system was subjected to per residues and pairwise residues decomposition to point residues that contribute majorly in system stabilization and lower energy. Molecular docking simulation studies demonstrated 64 residues from the TLR3 receptor that are in direct contact with the MEPVC but per residue decomposition assay illustrated that among the residues only Hie31, Phe55, Glu98, Hie100, Met102, Ile121, Thr122, Asp124, Glu146, Glu146, Glu174, Ser176, Phe198, Asn200, Ser225, Met249, Asp251, Tyr254, Tyr273, Phe275, Tyr278, Tyr297, Glu323, Hie324, and Tyr348 are hotspot as they contribute rigoursly in binding interaction with MEPVC at the docked side. The side chain of Hie100, Thr122, Asp124, Glu174, Ser176, Tyr254, and Glu323 contribute significantly in chemical interactions and have energy value in following order: −2.86602 kcal/mol, −3.71782 kcal/mol, −3.77019 kcal/mol, −3.80724 kcal/mol, −2.71475 kcal/mol, – 2.40187 kcal/mol, and −3.54158 kcal/mol, respectively. To these TLR3 hotspot residues, the MEPVC interacting residues were also observed in quite lower energies illustrating high affinity for the receptor residues for chemical interactions. From MEPVC, ASN633 (−4.11274 kcal/mol), TYR638 (2.83056 kcal/mol), VAL642 (−2.12862 kcal/mol), ARG643 (−6.59981 kcal/mol), GLU675 (−2.57531 kcal/mol), TRP682 (−2.54238 kcal/mol), ARG705 (−3.1088 kcal/mol), and ARG706 (−4.45088 kcal/mol) are favorable residues in stable complex formation. The hotspot residues for both TLR3 and MEPVC are shown in **S-Fig.2**.

### 3.15. Frame –wise Energy Decomposition

The binding free energy of the TLR3-MEPVC complex, TLR3 receptor, MEPVC and the net system energy is further decomposed into 100 frames extracted from simulation trajectories (**S-Fig.3**). This information deemed vital in predicting the simulation time where higher intermolecular affinity was observed and can guide about the most suitable docked conformation. In general the complex, receptor and construct energies are higher in PB compared to GB but for the total energy, the PB energies are quite lower for frames in contrast to GB. For the complex, the min, max and average binding energy reported are – 67264.7 kcal/mol, −66901.5 kcal/mol, and −67069.5 kcal/mol, respectively in GB. The PB max frame energy is −66751.4 kcal/mol, min is −67120.2 kcal/mol and average is −66921.6 kcal/mol. The GB receptor max is −57111.5 kcal/mol whereas the min is −57381.8 kcal/mol, and average is −57242 kcal/mol. For PB receptor, max of −56961 kcal/mol of energy was noticed compared to the min −57249 kcal/mol. The MEPVC is showing high average energy (−9773.16 kcal/mol in GB and −9742.47 kcal/mol in PB), max (−9672.42 kcal/mol in GB and – 9644.54 kcal/mol in PB) and min (−9838.37 kcal/mol in GB and −9811.3 kcal/mol in PB). The net average energy for frames in GB is −53.8024 kcal /mol (max, −33.978 kcal/mol and min, – 72.791 kcal/mol) and in PB is −89.0949 kcal/mol (max, −65.151 kcal/mol and −120.384 kcal/mol).

### 3.16. Pair-wise Energy Contribution

Pair-wise energy contribution to the net energy of the system was accomplished in order understand pair residues role from both TLR3 and MEPVC in complex stability. We found that the Thr122 and Asp124 (−4.56 kcal/mol in GB and −5.45 kcal/mol in PB), Glu174 and Ser176 (−3.45 kcal/mol in GB and −3.77 kcal/mol in PB), Glu323 and Hie324 (−2.86 kcal/mol in GB and −3.99 kcal/mol in PB) of TLR3 receptor have high combine contribution to the net energy. In case of MEPVC, Asn633 and Thr634 (−3.21 kcal/mol in both GB and PB), Val642 and Arg643 (−5,87 kcal/mol in GB and −3.27 kcal/mol in PB) and Arg705 and Arg708 (−2.74 kcal/mol and 2.04 kcal/mol).

## 4. Conclusions

Taken together, we characterized SARS-CoV-2 spike glycoprotein for antigenic peptides and proposed a MEPVC by means of several computational immunological methods and biophysical calculations. The outcomes of this study could save time and associated cost that go into experimental epitope targets study. The MEPVC is capable of activating all components of the host immune system, have suitable structural and physicochemical properties. Also, it seems to have very stable dynamics with TLR3 innate immune receptor and thus has higher chances of presentation to the host immune system. However, additional *in vivo* and *in vitro* experimentations are needed to disclose its potential in fight against COVID-19.

## 6. Acknowledgment

Authors are highly grateful to the Pakistan-United States Science and Technology Cooperation Program (Grant No. Pak-US/2017/360) for granting the financial assistance.

## 7. Supplementary File

**S-Fig.1**. PROSA-Z energy plot for the MEPVC.

**S-Fig.2**. Residue wise decomposition of net binding energy into TLR3 receptor and MEPVC interacting residues. Top (GB) and bottom (PB).

**S-Fig.3**. Binding energy decomposition per frame for TLR3-MEPVC complex (A), TLR3 receptor (B), MEPVC (C) and net energy (D).

**S-Table 1**. B cell epitopes predicted for the SARS-CoV-2 spike glycoprotein.

**S-Table 2**. Top 5 refined model of the MEPVC. The input MEPVC is also provided.

